# Phenotypic and transcriptomic responses to stress differ according to population geography in an invasive species

**DOI:** 10.1101/2020.09.02.279315

**Authors:** Pierre Marin, Angelo Jaquet, Justine Picarle, Marie Fablet, Vincent Merel, Marie-Laure Delignette-Muller, Mariana Galvão Ferrarini, Patricia Gibert, Cristina Vieira

**Author notes:** Corresponding author &.

## Abstract

**Background:** Adaptation to rapid environmental changes must occur within a short time scale. In this context, studies of invasive species may provide insights into the underlying mechanisms of rapid adaptation as these species have repeatedly encountered and successfully adapted to novel environmental conditions. Here we investigated how invasive and non-invasive populations of *D. suzukii* deal with an oxidative stress at both the phenotypic and molecular level. We also investigated the impact of transposable element insertions on the differential gene expression between genotypes in response to oxidative stress.

**Results:** Invasive populations lived longer in the untreated condition than non-invasive Japanese populations. As expected, lifespan was greatly reduced following exposure to paraquat, but this reduction varied among genotypes (a genotype by environment interaction, GEI) with invasive genotypes appearing more affected by exposure than non-invasive genotypes. We also performed transcriptomic sequencing of selected genotypes upon and without paraquat and detected a large number of genes differentially expressed, distinguishing the genotypes in the untreated environment. While a small core set of genes were differentially expressed by all genotypes following paraquat exposure, much of the response of each population was unique. Interestingly, we identified a set of genes presenting genotype by environment interaction (GEI). Many of these differences may reflect signatures of history of past adaptation. Transposable elements (TEs) were not activated after oxidative stress and differentially expressed (DE) genes were significantly depleted of TEs.

**Conclusion:** In the decade since the invasion from the south of Asia, invasive populations of *D. suzukii* have diverged from populations in the native area regarding their genetic response to oxidative stress. This suggests that such transcriptomic changes could be involved in the rapid adaptation to local environments.

## Introduction

Rapid environmental changes, particularly related to human activity, can decisively affect living organisms, who must respond to them within a short-time scale. Understanding the mechanisms underlying these rapid responses is challenging and could help predict organism and species survival in the face of global environmental changes. The rapid adaptation of invasive species to new environments, some quite different than ancestral environments, may provide insights into such mechanisms [1,2] including hormonal regulation of suites of traits, or epigenetic gene regulation [3–6]. Phenotypic plasticity, *i*.*e*., the ability of a genotype to express different phenotypes in different environments, is a possible explanation to the success of invasive species, particularly in the case of founder populations depleted of genetic variations [3–5,7].

Genetic diversity could rapidly increase following environmental stress if there is an activation of transposable elements (TEs) or if the epigenetic control is disturbed. TEs, which are repeated sequences that can move around genomes, were discovered by B. McClintock in the 50’ [8]. Depending on where TEs are inserted within the genome, they can affect the fitness of its host organism. The vast majority of new TE insertions are neutral or deleterious, and purifying selection is expected to remove them or favours their silencing [9–11]. However, some TE insertions may be advantageous and facilitate adaptation in different environments [6,12–22]. Such adaptive effects have been previously observed in response to both biotic (*e*.*g*., virus infection) and abiotic (*e*.*g*., oxidative stress) stress [19,20]. Moreover, stress-induced changes in the epigenetic regulation of TEs, which is often sensitive to environmental cues [14,21,23], has already been described to rapidly generate potentially advantageous changes in nearby gene regulation and facilitate rapid adaptation to environmental stress [9,13].

Here, we examined variation in the oxidative stress response of invasive and non-invasive populations of *Drosophila suzukii* with a focus on molecular mechanisms potentially underlying the observed phenotypic differences. *D. suzukii* is an Asian species of the *melanogaster* group that invaded North America and Europe in 2008 [24–28]. Outside of Asia, *D. suzukii* is now found in both North and South America, and throughout most of Europe, from southern Spain easterly into Poland, Ukraine and Russia [25–28]. As *D. suzukii* has spread throughout the world, it has encountered and successfully colonised many different, potentially stressful environments.

Paraquat (*N,N′-dimethyl-4,4′-bipyridinium dichloride*) is one of the most widely used herbicide in the world [29,30]. Exposure to paraquat leads to the production of ROS (reactive oxygen species) and has often been used in the lab as a proxy to study oxidative stress [31–34]. Resistance to oxidative stress has been associated with extended lifespan [34–36], a trait possibly under selection during invasion of a new area. Furthermore, paraquat has been banned since 2007 in Europe but is still used in the U.S.A and Japan.

In this study, we compared field-sampled *D. suzukii* genotypes collected in their native area of Japan with genotypes collected in invaded areas in the U.S.A and France. For each genotype, we measured lifespan in both the presence and absence of paraquat, where we identified an effect of genotype and a genotype-by-environment interaction effect (GEI). We went further by examining the transcriptomic response of single genotypes from each location along with analysis of TE expression. We found substantial differences among genotypes in patterns of gene expression related to oxidative stress that may explain the observed phenotypic differences and reflect population history. This work highlights the local adaptation to environmental conditions of the genotypes within a short-time scale.

## Results

### Among population variation for lifespan and oxidative resistance

We have measured lifespan and oxidative stress in a total of 27 isofemale lines from six geographical regions coming from the U.S.A, France and Japan. As expected, oxidative stress had a strong negative effect on survival, with an average decrease in lifespan of 80% when paraquat was present in the medium (multiplicative coefficient of 0.20, Fig. 1). Median lifespans of flies are presented in Table 1 for each population, sex and treatment and statistical analysis of survival is presented in Fig. 1 and Table S1. Sex differences in lifespan and changes in lifespan in response to stress are present in some species [37]. However, we did not find a main effect of sex or any significant interactions with sex in our preliminary statistical model (see materials and methods). Therefore, male and female data were pooled for subsequent analysis.

**Table 1.** Median lifespan (days) by sex, treatment, and population with bracketed 0.25 & 0.75 quantiles. Values were calculated from estimated median (see M&M) at population level (line effects into the populations were estimated as a random effect from the linear mixed model).

|  | Sapporo<br>(Japan) | Tokyo<br>(Japan) | Montpellier<br>(France) | Paris<br>(France) | Dayton<br>(U.S.A) | Watsonville<br>(U.S.A) |
| --- | --- | --- | --- | --- | --- | --- |
| Females control | 32.0 (26.0-39.8) | 32.0 (29.6-41.0) | 40.5 (35.3-46.3) | 41.0 (34.5-49.3) | 51.4 (39.7-54.6) | 34.8 (27.4-42.4) |
| Females paraquat | 6.8 (5.0-9.1) | 5.3 (5.0-7.7) | 5.5 (5.2-6.3) | 7.1 (5.4-8.2) | 9.1 (8.7-9.8) | 5.1 (3.8-6.5) |
| Males control | 32.6 (24.65-37.85) | 33.2 (27.32-44.43) | 39.4 (36.69-44.41) | 39.9 (36.48-48.3) | 40.4 (39.46-46.96) | 33.9 (26.51-38.92) |
| Males paraquat | 6.1 (4.85-7.57) | 7.0 (5.89-8.2) | 7.5 (6.66-8.08) | 7.6 (6.81-9.02) | 12.0 (10.78-12.91) | 4.4 (3.16-5.17) |

**Fig. 1.**
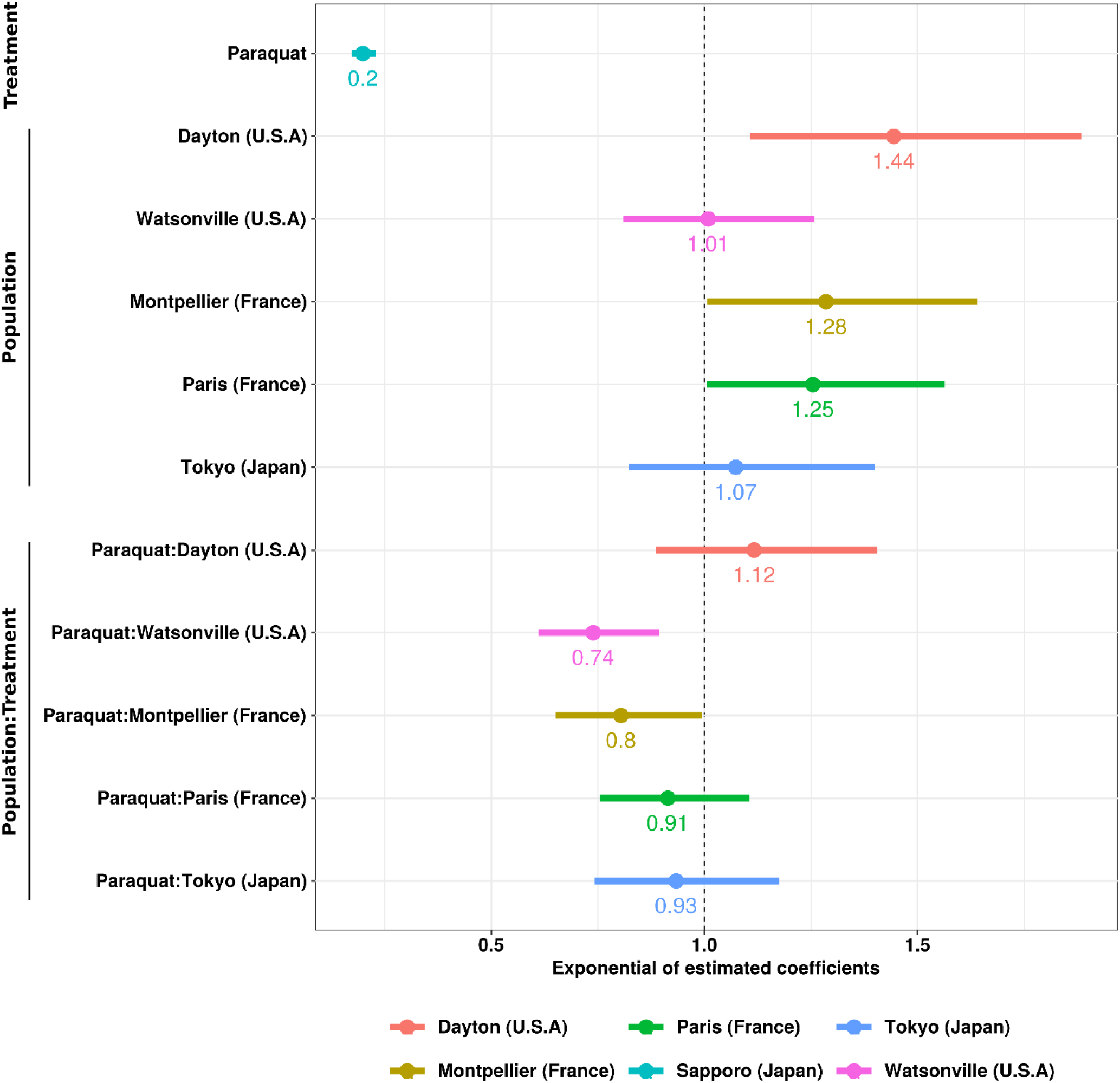
Lifespans under control and paraquat-treated conditions relative to the Japan Sapporo population with confidence intervals (0.95). The relative values are within each treatment group and can be interpreted as a multiplicator effect compared to the reference level (Sapporo population, female and control condition). The intercept, which is the basal level of the Sapporo females in untreated condition, is equal to 31.96 days. Paraquat correspond to the mean effect of the treatment on Sapporo. Mean values of the populations correspond to the effect of the population on the lifespan compared to Sapporo in untreated condition and the last are the interaction term after paraquat exposure. As examples, the effect of paraquat reduced the lifespan of Sapporo to 20% (0.2) of the initial value in untreated condition. Values higher than 1 indicate an increase in the lifespan compared to Sapporo, while below 1 this indicate a higher sensitivity (*e*.*g*., Paraquat:Watsonville correspond to the interaction term and indicates that after paraquat exposure, Watsonville remains more sensitive than Sapporo with a difference of 26% (1-0.74)).

In the untreated condition, flies from the two Japanese populations had the shortest lifespan and were not significantly different. For flies sampled in the United States, those from Watsonville had a median lifespan very similar to the Japanese populations and were not different from the reference Sapporo population (Fig. 1, value = 1.01, corresponding to about 1% greater lifespan than the reference Sapporo population). However, flies from Dayton lived the longest (value=1.44, a 44% relative increase). The two populations collected in France lived on average 25-28% longer than flies in the Sapporo population (1.25 and 1.28 for Paris and Montpellier, respectively).

The decline in lifespan following paraquat treatment was variable among populations (genotype by environment interaction). Compared to the Sapporo reference population, there were non-significant reductions in resistance for Tokyo, Paris and Dayton. Populations from Watsonville and Montpellier were significantly more sensitive to paraquat treatment, with reductions in lifespan of 14.8% and 16% respectively (multiplicative effects of 0.20*0.74 and 0.20*0.80 in Fig. 1). We observed a low but significant correlation among genotypes for lifespan across the two environments (r= 0.28, p-value = 3.3E^-4^, Fig. S1).

### Transcriptomic variability among genotypes

We quantified gene expression of three genotypes in somatic tissues, one from each geographical sampling location (Montpellier (MT47): France, Watsonville (W120): U.S.A & Sapporo (S29): Japan), hereafter referred by the country where flies were sampled. We choose these three genotypes because of their difference in lifespan.

A Principal Component Analysis (PCA) of gene read counts (Fig. 2) clearly showed genotype-specific clustering, independent of the treatment. To evaluate variation in the transcriptomic response of each genotype to paraquat treatment, we computed the coefficient of variation (CV) for each differentially expressed (DE) genes between control and treated flies (Fig. S2). CV distributions were significantly different across genotypes (paired Wilcoxon test, p-values < 0,01), which suggests significant genotype by environment interaction for transcriptomic response. The number of differentially expressed (DE) genes identified, (i) in pairwise comparisons between genotypes in control conditions, (ii) in comparisons between untreated and oxidative stress conditions for each genotype, and (iii) in pairwise comparisons between genotypes following paraquat treatment are presented in Table 2. The distribution and values of the CV were in agreement with the distribution of DE genes shown in the Table 2, suggesting that the difference in DE gene proportions between the genotypes are due to biological variation and not a bias of statistical power.

**Table 2.** Number of DE genes between genotypes and treatments. Pairwise comparisons between (A) untreated genotypes, (B) between treated and untreated flies within each genotype, and (C) in pairwise comparisons of paraquat-treated flies between different genotypes. The threshold for identifying DE genes was an adjusted p-value ≤ 0.01 and absolute log_2_fold-change ≥ 1. The proportion of DE genes is the percentage of DE genes in the expressed transcriptome (14538).

|  | Contrast | DE genes | Up-regulated | Down-regulated | DE proportion (%) |
| --- | --- | --- | --- | --- | --- |
| A | France\|Japan control | 524 | 175 | 349 | 3.6 |
|  | France\|U.S.A control | 715 | 471 | 244 | 4.92 |
|  | U.S.A\|Japan control | 1023 | 208 | 815 | 7.04 |
| B | Japan (paraquat\|control) | 122 | 74 | 48 | 0.84 |
|  | France (paraquat\|control) | 531 | 354 | 177 | 3.65 |
|  | U.S.A (paraquat\|control) | 281 | 214 | 67 | 1.93 |
| C | France\|Japan paraquat | 138 | 105 | 33 | 0.95 |
|  | France\|U.S.A paraquat | 65 | 19 | 46 | 0.45 |
|  | U.S.A\|Japan paraquat | 62 | 57 | 5 | 0.43 |

**Fig. 2.**
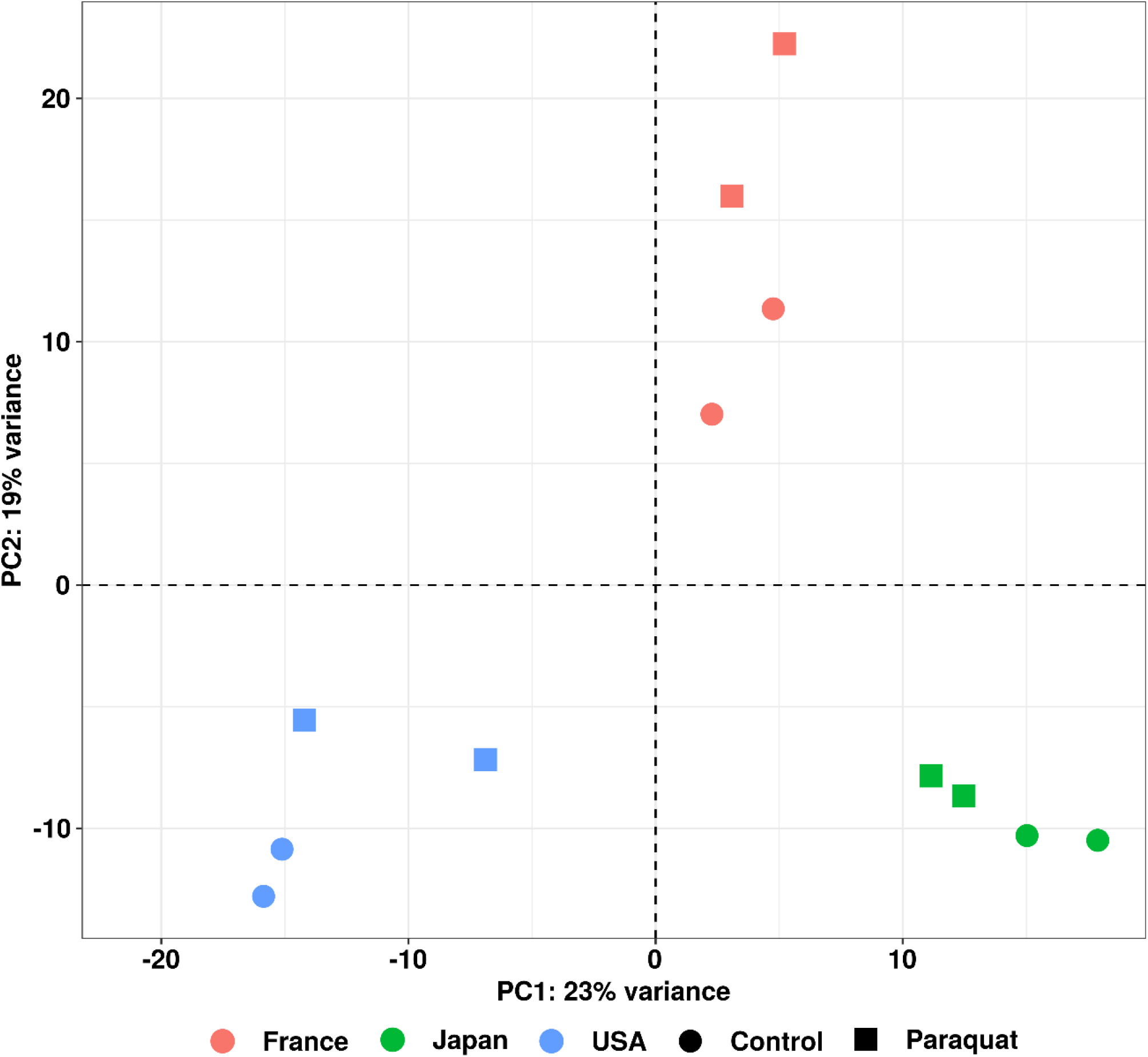
PCA analysis using normalized read counts from DESeq2. Dots correspond to the biological samples with the U.S.A in blue, Japan in red and France in green. Circles and squares correspond respectively to control and paraquat treatment.

**Table 3.** Insertion position of TEs in genes differentially expressed.

### Genotypic variation in gene expression in untreated flies

Pairwise comparisons of gene expression of untreated flies between the three genotypes revealed 715 DE genes between France and U.S.A (4.92% of the total transcriptome), 524 between France and Japan (3.6%), and 1023 between U.S.A and Japan (7.04%) (Table 2 and Fig. S3). Most of these DE genes (∼70%) had an absolute log_2_fold change below 2 (Fig. S3) and only 60 had an absolute log_2_fold change higher than 5.

To further examine these DE genes, we performed a Gene Ontology analysis (Fig. 3). The rationale was to identify transcriptomic differences possibly related to adaptation of the different genotypes to their respective environments. In the comparison of France vs U.S.A, there were fewer enriched terms (all of them from up-regulated genes in France) when compared to France *vs* Japan or U.S.A *vs* Japan. In the comparisons of France *vs* U.S.A, enriched terms came from down-regulated genes in the Japan genotype. These results suggest a greater similarity between the two invasive genotypes, France and U.S.A. The greater enrichment of GO terms in comparisons between Japan and either the U.S.A or France suggests this population is extremely different than the other two.

**Fig. 3.**
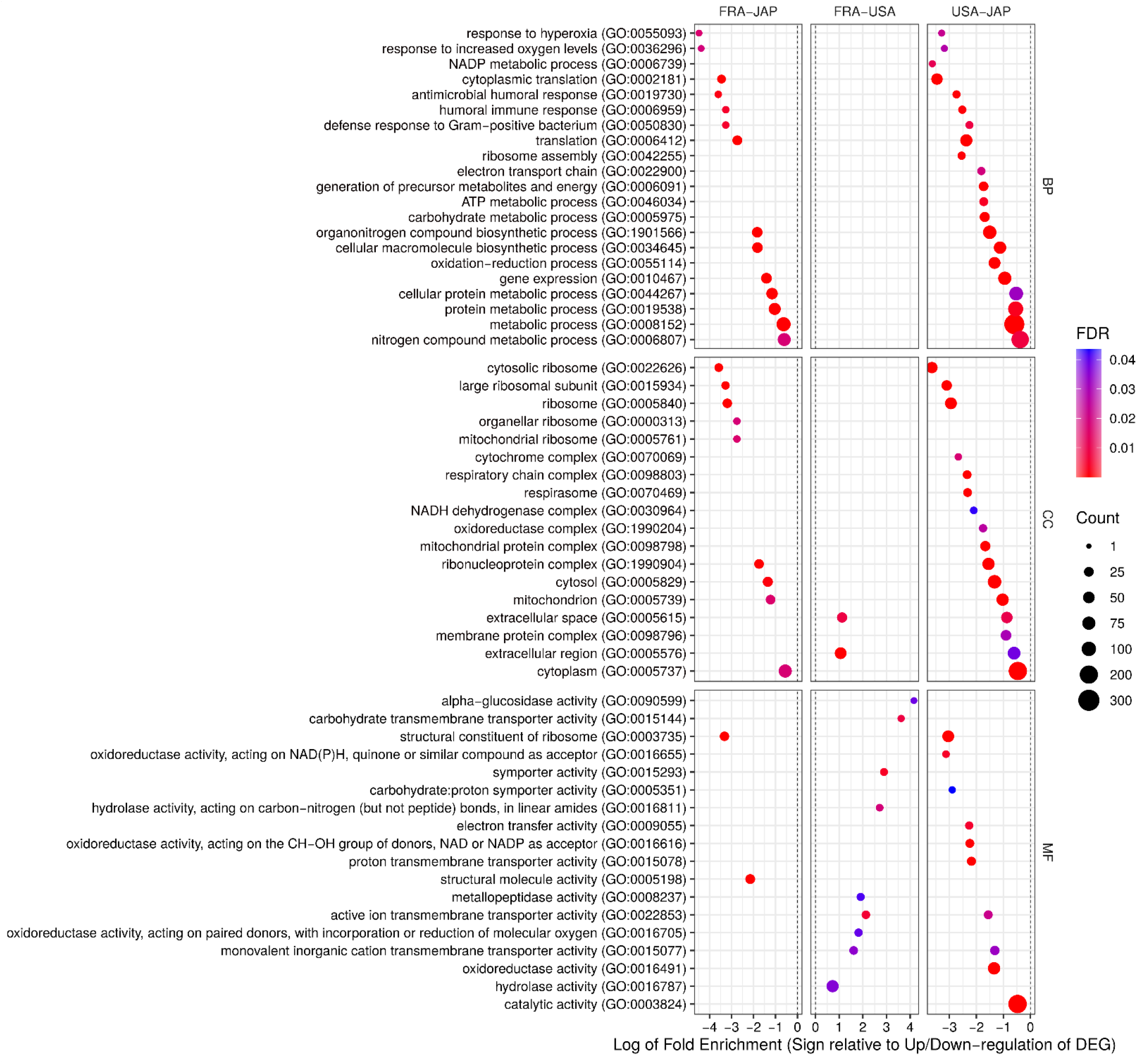
Gene ontology enrichment analysis for all genotypes in control condition. BP: Biological Process; CC: Cellular Components; MF: Molecular Function. The size of dots corresponds to the number of genes in each category and the colour to the FDR. Gene ratio correspond to the number of genes from our data compared to genes within a GO term. Down-regulated genes from pairwise comparisons (*e*.*g*., France-Japan) are symbolized with negative gene ratio and a positive gene ratio for up-regulated genes.

We detected 44 GO terms shared between the invasive genotypes (France and U.S.A) in comparison with the non-invasive Japan genotype. These terms were mainly related to translation, protein metabolic process, ribosome biogenesis, response to hyperoxia, and immune response (antibacterial related). All of these terms were down-regulated in the invasive genotypes (U.S.A or France) when compared to the non-invasive Japan genotype. We also detected other functional terms in molecular function (MF) that seemed to be specifically down-regulated in the U.S.A genotype (so they appear in both U.S.A *vs* Japan and France *vs* U.S.A results): carbohydrate transport and energy metabolism. It is plausible to say that these functions are compromised in the U.S.A genotype.

Taken together, these enrichment analyses suggest transcriptomic differences in translation, protein metabolic process, ribosome biogenesis, response to hyperoxia, and immune response (antibacterial related), which have been down-regulated in invasive genotypes compared to the non-invasive Japanese genotype.

### Oxidative stress induces genes upregulation in invasive genotypes

We compared changes in gene expression between flies in control and oxidative conditions and identified a total of 659 unique DE genes across the 3 genotypes (Fig. 4 and Table 2). The Japan genotype had the fewest DE genes (122 genes, representing 1.10% of the transcriptome) in response to paraquat treatment, followed by the U.S.A (281 genes, 2.46%) and France (531 genes, 4.51%). Of all DE genes, most were upregulated upon oxidative stress (435/659). When comparing DE genes among genotypes, we observed that fewer genes were shared between Japan and the other two genotypes (Fig. 4), with respectively 4 and 23 genes uniquely shared with U.S.A and France. The comparison between France and U.S.A showed that a greater number of DE genes were uniquely shared (114) between these two genotypes.

**Fig. 4.**
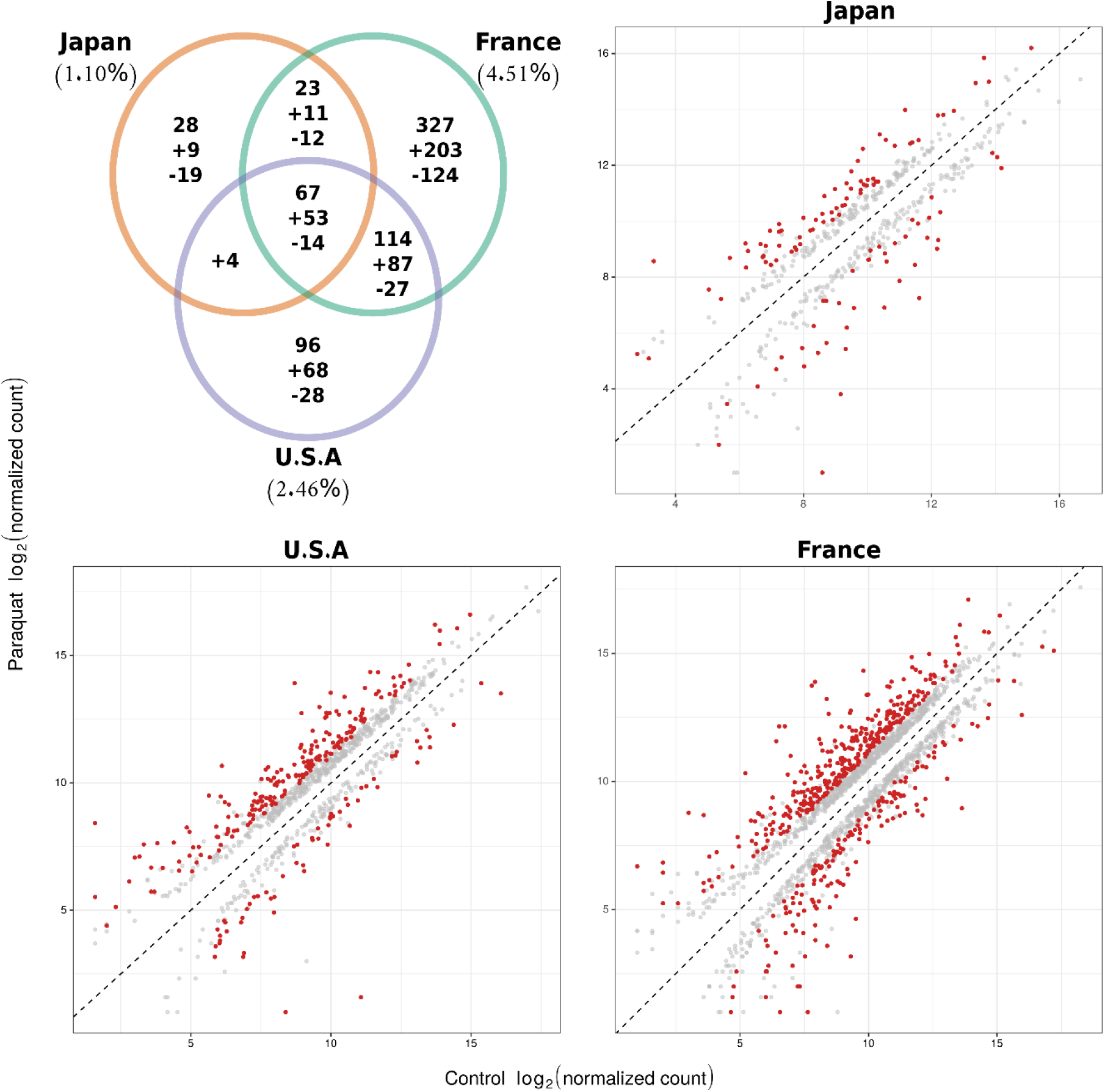
Gene expression between control and paraquat conditions. A Venn diagram of shared and unique DE genes identified in comparisons of paraquat-treated and control flies within each population after paraquat exposure (top left). Scatter plots of log_2_ normalized read count for Japan (top right), U.S.A and France (bottom panels) comparing control and treated flies. Individual genes are indicated by dots. Red colour corresponds to significant DE genes (see materials and methods).

A gene ontology enrichment analysis for each genotype was performed with 621 annotated genes out of the 659 DE genes. We were able to detect enriched terms for down-regulated genes in the Japan genotype and for up-regulated genes in the U.S.A and France genotypes. These observations are in accordance with the fact that, a functional major up-regulation of genes in response to paraquat was only observed in invasive genotypes. When comparing the GO terms enriched in up-regulated genes from invasive genotypes (Fig. 5), terms such as ligase activity, oxidation-reduction, ATP binding, drug binding and ion binding were common to France and U.S.A. As observed in related species, paraquat can indeed cause DNA damage via oxidative stress [31]. The French genotype had a greater number of specific enriched terms, mostly related to DNA repair (including aforementioned ligase activity and telomere maintenance, among others), protein translation, protein refolding and mitochondrion. The U.S.A genotype had other enriched terms related to carbohydrate metabolism, detoxification, and response to metal ion. There were no enriched GO terms among up-regulated genes in the Japan genotype. Enriched terms for down-regulated genes in the Japan genotype were mainly related to immune response (to bacteria), response to increased oxygen levels (hyperoxia) and peptidase activity (Fig. 5). Overall, it appeared that while paraquat induced increased expression for genes related to oxidation-reduction, detoxification, drug/metal binding, DNA repair and protein refolding in invasive genotypes, it reduced the expression of important genes for the antioxidant response in the non-invasive genotype.

**Fig. 5.**
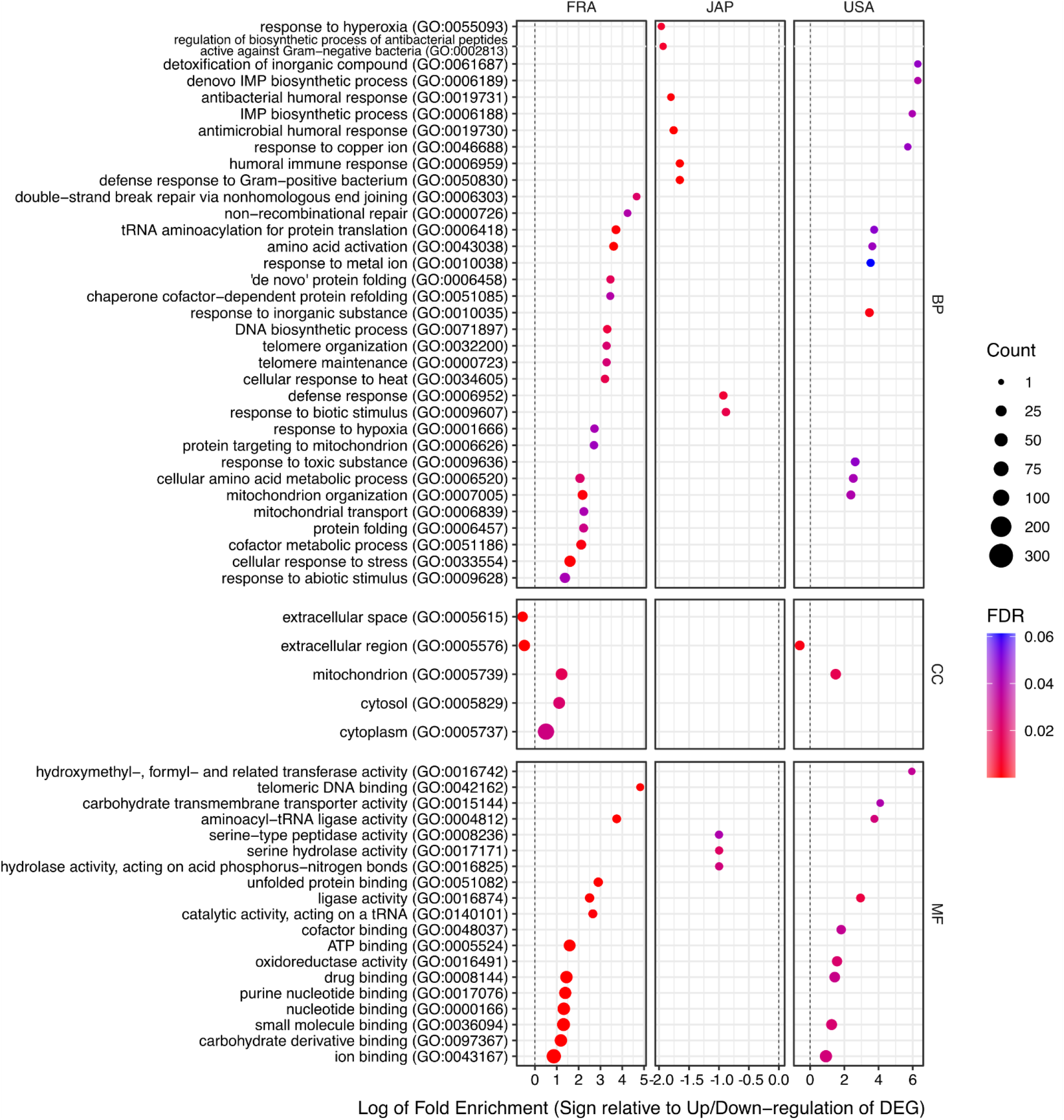
Gene Ontology analysis for up- and down-regulated genes induced upon treatment with paraquat. Up- and downregulated DE gene lists from the three pairwise comparisons (Paraquat vs Control) were used in this analysis in order to detect enriched functions. For this we used: 243 genes up-regulated and 105 genes downregulated in the French strain, 134 genes up-regulated and 42 genes down-regulated in the USA strain and 31 genes down-regulated from the Japanese strain. No functional enrichment with up-regulated genes from the Japanese genotype was detected.

### DE genes common to the three genotypes were mostly upregulated with oxidative stress

From a total of 659 unique DE genes between control and paraquat exposure, 67 were shared by all genotypes. This set of core genes were regulated in the same way for the three genotypes: 14 down-regulated (from log_2_FC = −1.03 to −10.8) and 53 upregulated (from log_2_FC = 1.03 to 10.48) (Fig. 6).

**Fig. 6.**
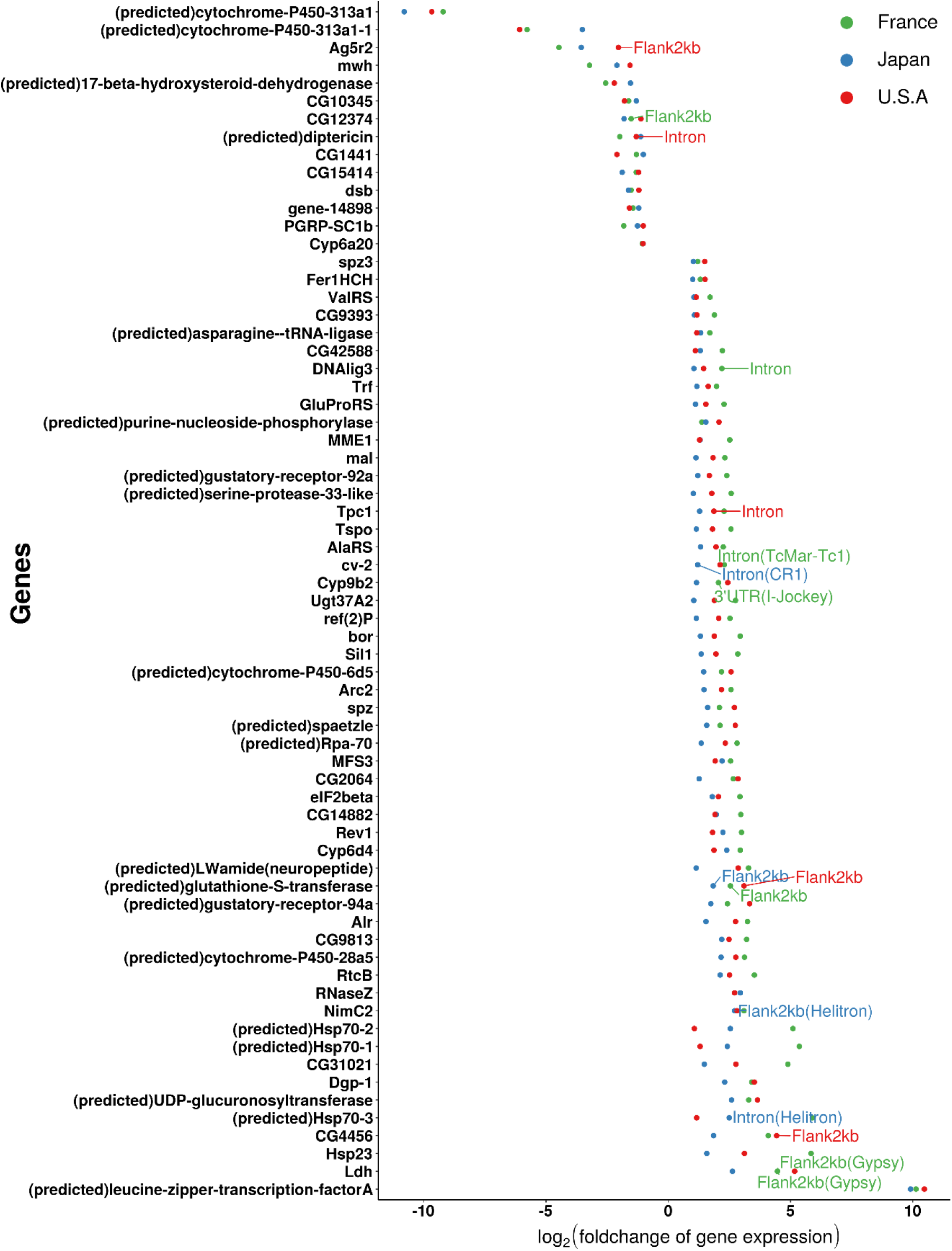
DE genes (67) shared by all genotypes after paraquat exposure (U.S.A in red, France in green and Japan in blue). The plotted labels refer to a TE insertion (frequency ≥ 0.5) within a gene (intron, exon, 5’ or 3’UTR) or within the 2kb flanking regions of a gene.

Among those up-regulated following paraquat treatment, we found genes related to stress response such as *Hsp* and *Cyp* genes families. The strongest up-regulated genes were a predicted gene encoding for a transcription factor A (log_2_fold-change = 10) and other genes in the *Hsp* gene family. Among the strongest down-regulated genes, we identified a cytochrome P450 gene that was the top down-regulated gene in all 3 genotypes and has a log_2_FC < −10 in the Japanese genotype.

We performed a GO enrichment analysis for the set of 67 genes common to the three genotypes. For down-regulated genes, only 9 of the 14 genes had a homolog in *D. melanogaster*. Enriched terms were associated with peptidoglycan metabolic process and negative regulation of NK cell differentiation involved in the immune response. However, all enriched terms were related to two genes: *PGRP-SC1a* and *PGRP-SC1b. PGRPs* (Peptidoglycan recognition proteins) are important in recognizing and degrading bacterial peptidoglycan, although *PGRP-SC1b* has not shown antibacterial activity and may instead be a scavenger protein [38]. Out of 53 up-regulated genes, 37 had homologs in *D. melanogaster*. Enrichment analysis on this set of genes identified only one significant GO term: ligase activity (which is related to DNA repair). Four of the five genes within this GO term were tRNA-ligases, which may play a role in protecting cells against oxidative damage following their translocation into the nucleus [39].

### The stress response is variable across genotypes

We identified a total of 213 unique genes with a significant GEI, which represent the set of genes with expression differentially modulated by oxidative stress according to genotype (Fig. 7). When comparing differences in the response of invasive genotypes to the non-invasive Japan genotype, we found 62 differentially modulated genes with the U.S.A genotype and 138 with the France genotype (Table 2). Most of these differences were due to greater up-regulation DE genes in the invasive genotypes (57/62 and 105/138). We identified 52 genes where the GEI was driven by a differential response in only one genotype compared to the other two. This included 22 genes differentially modulated in the France genotype compared to Japan and U.S.A, 14 in Japan compare to France and U.S.A, and 16 for U.S.A against others. We have presented some examples of these genes (Fig. 7 and Table S2), selected for the greatest log_2_fold change and illustrating cases in which the magnitude of the response to paraquat differed among genotypes. For example, *dysc* and *FarO* were down-regulated in France and upregulated in U.S.A and Japan. The *Hsp* genes *Hsp68* and *Hsp70Aa* were strongly up-regulated following paraquat treatment in France, with a log_2_FC ≥ 2, compared to the much-reduced changes in expression in Japan or U.S.A. In the USA genotype, *Mec2* was strongly down-regulated compared to the increased expression following treatment in the other genotypes. Oxidative stress appeared to upregulate *CCHa2, RpL40* and *Tsf1* only for the Japanese genotype. These examples highlight the potential effect of genotype-specific responses to oxidative stress.

**Fig. 7.**
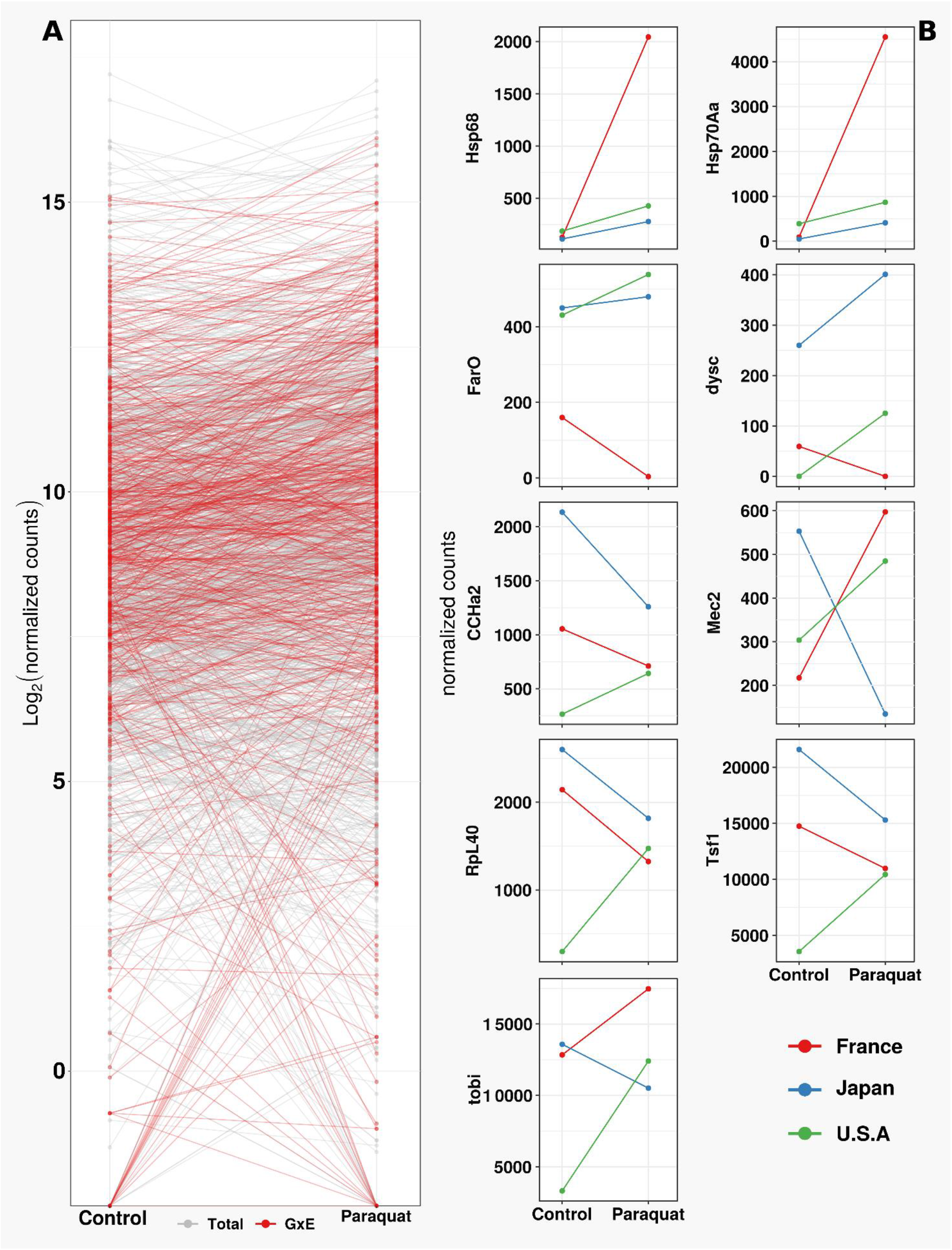
Reaction norms between control and paraquat for DE genes using normalized log_2_ read counts. Reaction norms of all DE genes, with red for the GEI ones, while grey are DE genes without GEI. Examples of 9 DE genes with GEI, with colours referred to the genotype (red for France, blue for Japan and green for the U.S.A)

### TE expression is not sensitive to oxidative stress

Environmental changes can affect the expression of TEs by lifting epigenetic repressive regulation mechanisms [17]. In our experiments, TE expression levels were very low, and reads corresponding to TEs did not exceed 3.8 to 7.1% of the total transcriptome. In control condition, differentially expressed TEs (DETEs) identified in pairwise comparisons between genotypes represented from 3.08 (48 families) to 5.91% (92) of total number of TE families annotated in the *D. suzukii* genome (Table S3). The U.S.A genotype exhibited a greater level of DE of TEs compared to the French or Japanese genotypes, with almost 70 TE families up-regulated in U.S.A genotype in comparisons with either France or Japan. Moreover, a similar number of both up- and down-regulated DETES were identified in the comparison between France and Japan (Fig. 8).

**Fig. 8.**
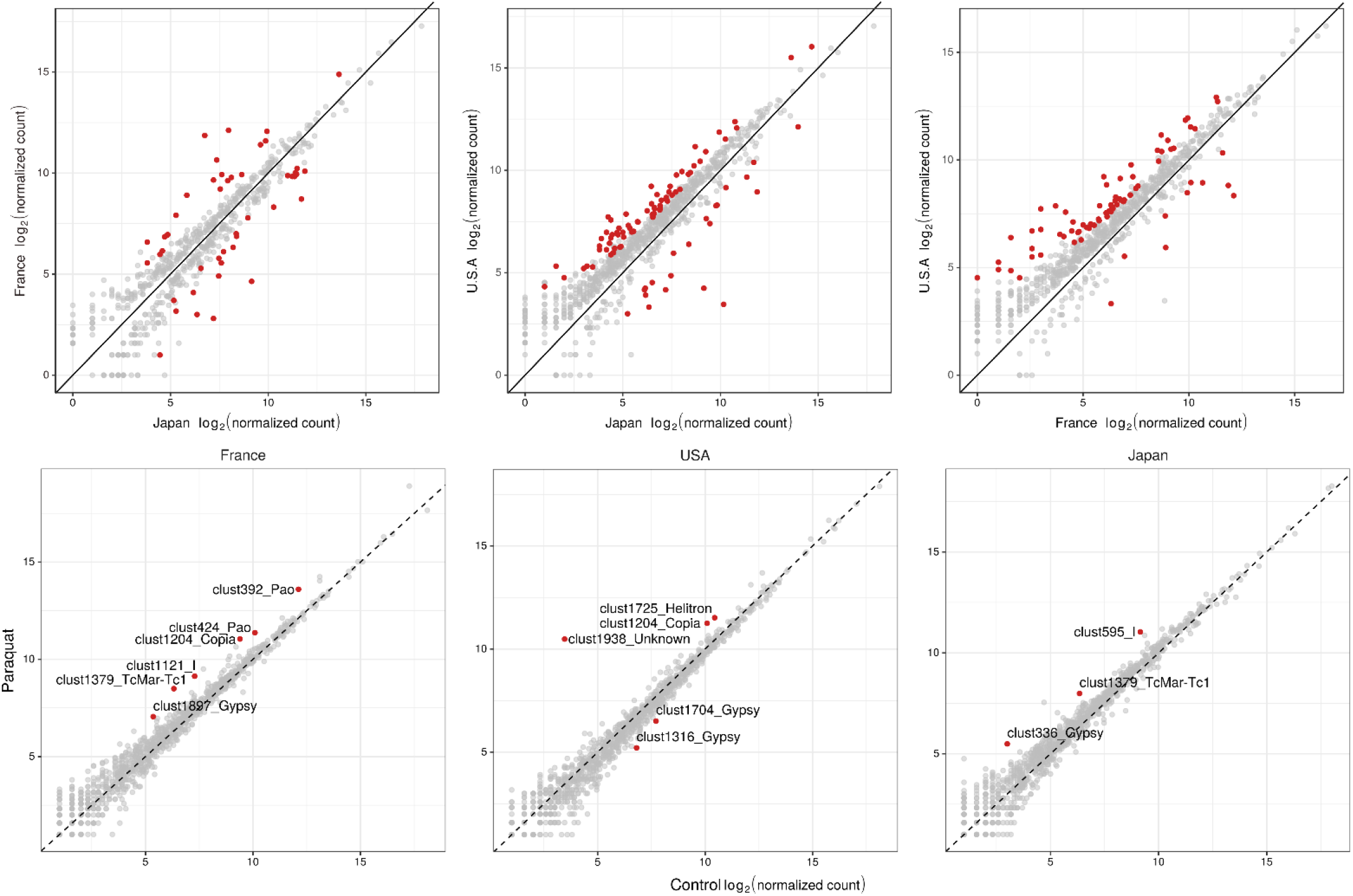
TE expression between genotypes (upper panel) and between control and paraquat conditions for each genotype (bottom panel). Scatter plots represent the log_2_ normalized read counts. Individual TE are indicated by dots. Red colour corresponds to significant DE TEs (see M&M).

After paraquat exposure, very few TE families changed in their expression levels (Table S3). In total, only 12 TE families were differentially expressed (Fig. 8). Six TE families in France and three in Japan were up-regulated. In the U.S.A genotype, differential expression of five TE families was observed, with three showing up-regulation and the remainder down-regulated. Among the DETEs, all classes of TE families were represented. We observed a differential expression in two of the genotypes in a Copia cluster and a Tc1 mariner cluster, which could suggest specific activation of these TE families upon oxidative stress.

### DE genes during oxidative stress are not enriched in TE insertions

TEs represent ∼33% of the *D. suzukii* genome and can potentially interfere with gene expression during stress [40]. To test for an enrichment or depletion of TE insertions around DE genes, we first test if the distribution of TEs in the three genomes was not significantly different (Chi-square test = 0.67, Table S4). We then tested the dependence of TE insertions and gene expression states (DE or not) after paraquat exposure (Table S5). Chi-square tests for the three genotypes showed that DE genes had fewer than expected TE insertions in genes and the 2kb flanking region (p-value < 0.05).

We then focused on all the 115 TE insertions present in the DE genes, the majority of which were in introns (57) or in ±2kb flanking regions (50) around DE genes (Table S6). Of the remaining 8 TEs, 7 were associated with up regulated genes (*JMJD4* (5’UTR), *Act42A* (exon), *Cyp9b2* (3’UTR), *CG8728* (3’UTR), *Cyp6a22* (3’UTR), *CG6834* (3’UTR), and one non annotated gene (exon). One insertion was associated with a down-regulated gene, *CG4409* (*exon*).

### Shared DE genes are not enriched with TE insertions

In agreement with a depletion of TE in DE genes, of the 67 shared DE genes that responded to paraquat treatment in similar ways across the three genotypes (Fig. 6), we founded 11 genes with one or more TE insertions. Among these 11 genes, only one (a gene predicted to encode a glutathione transferase) had a shared element present at the same position in all three genotypes (helitron family ∼1kb upstream the gene).

### Distribution of TEs among GEI genes

A GEI interaction indicates that the magnitude or direction of changes in expression following treatment could differ depending on the genotype. We found a total of 53 genes with at least one TE insertion. The DE genes showing evidence of a GEI, present the same distribution of TE insertion as all genes, except for genes with a GEI between France and U.S.A (p-value = 0.016, Table S5) in which they are less frequent and no insertion was shared in all genotypes. Also, the TE insertions were associated with high or low level of expression (Fig. 9 summarizes detected TE insertions in GEI DE genes for the different genotypes, also Table S7). For example, three genes were differentially expressed between France and the others genotypes; *FarO* (*Fatty acyl-CoA reductase*), *kelch* (which plays an essential role in oogenesis, where it is required for cytoskeletal organization), and *Hsp70-Aa* (a protein involved in response to heat shock and hypoxia). *Kelch* and *FarO* both had a TE insertion in France, with, respectively, a greater and lesser expression compared to other genotypes. *Hsp70-Aa* had an insertion in Japan and U.S.A and showed lower expression than in France. Another example is the gene *CG12520*, which has a TE insertion in the 3’UTR in Japan and a lower expression.

**Fig. 9.**
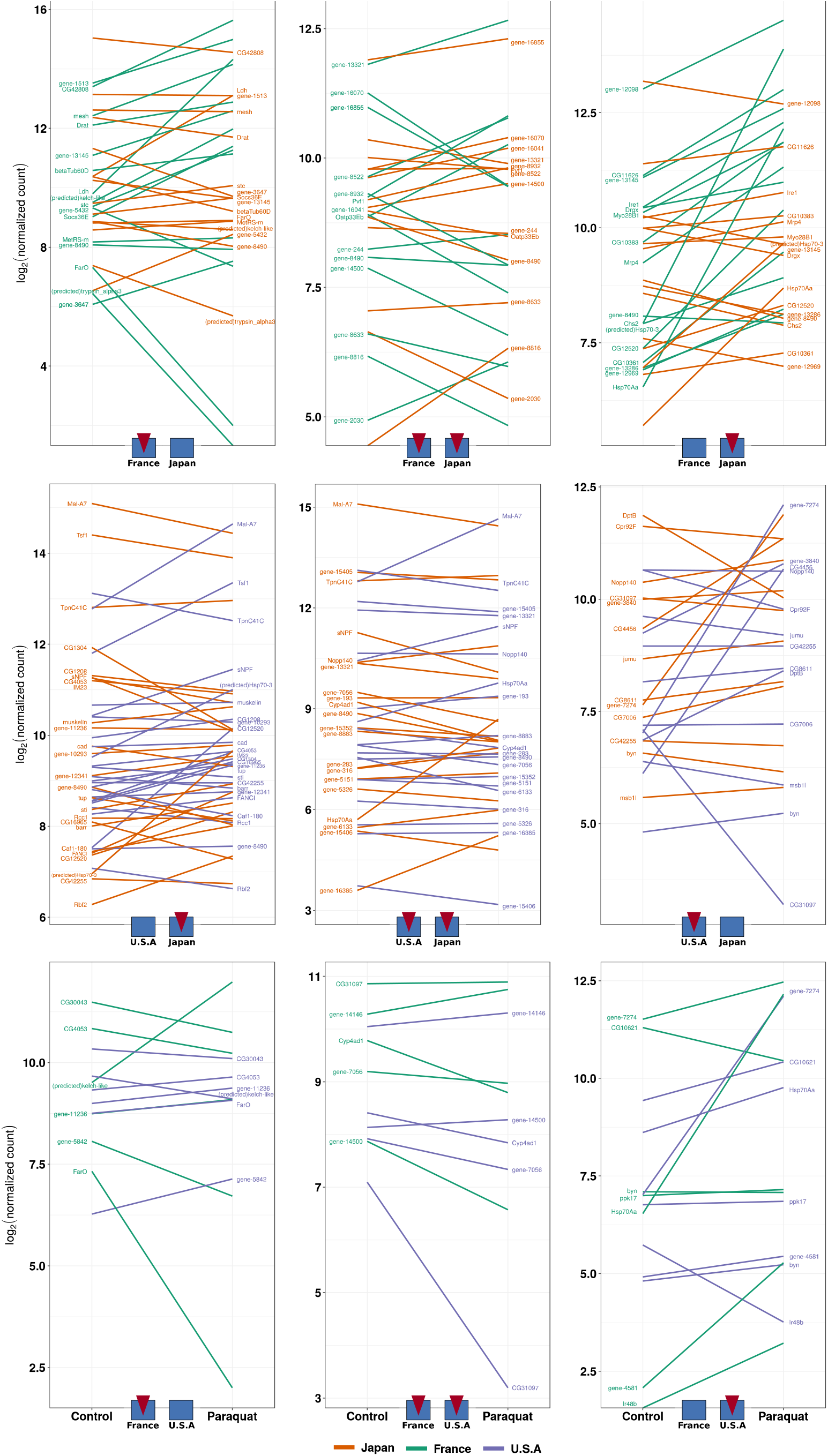
Expression differences (expressed as log_2_-fold-change) of paraquat-exposed flies of DE genes showing evidence of GEI with detected inserted transposable elements. Colour indicates genotypes with insertions that are either shared or unique among the populations (France and Japan on the top, U.S.A and Japan on the middle, France and U.S.A on the bottom).

TEs may impact neighbouring gene expression due to the transcription factor binding sites (TFBSs) they harbour. We investigate whether this was the case in our data set and focused on three TFBSs of the antioxidant responses elements group (ARE) [41]. We first analysed enrichment in TFBS for all TE families identified in the GEI genes (Fig. S4A). Of 196 TE families, 36 had at least one TFBS, most of them related to a CnC (Cap’n’collar) element. The TE sequences with putative TFBS were related to Pao and Gypsy families. We founded 4 GEI genes with TFBS linked to a TE insertion (Fig. S4B). Only one was annotated as *stc* gene (*shuttle craft*), which encodes a NFX1 family transcription factor implicated in modulating adult lifespan and aging [42].

## Discussion

### *D. suzukii* genotypes vary in lifespan and response to oxidative stress

Previous studies founded a positive association between stress resistance and extended lifespan or aging in *D. melanogaster* [34,43,44]. In this species, the ROS defences are mediated by both immune and antioxidant response pathways. A similar association may be expected in *D. suzukii*, a species which diverged from the melanogaster group ∼8 Mya. However, until now, no extensive study had been performed using *D. suzukii* wild-type genotypes. Here, we observed a significant positive correlation between lifespan in standard conditions and under oxidative stress.

However, not all fly genotypes responded to oxidative stress in the same way, resulting in a significant genotype-by-environment interaction (Fig. 1 and Table 1), in accordance to what was reported in *D. melanogaster* [32]. For example, Japanese populations had the lowest lifespan in the untreated condition but were more resistant to oxidative stress than genotypes from Watsonville (U.S.A) or Montpellier (France). This GEI suggests possible local adaptation of the different populations to paraquat, perhaps associated with differences in herbicide use in the three countries. Paraquat is one of the most used herbicides in the world and is widely used in Japan and U.S.A, but forbidden in Europe since 2007 [29,30]. The presence of *D. suzukii* in Europe has been reported since 2008, and flies are therefore unlikely to have encountered paraquat in the field since their arrival [24,27,45]. This could have resulted in a relaxed selective pressure for oxidative stress resistance and explain why the French Montpellier population was more sensitive than Japanese and American lines (except for Watsonville). The Paris population, on the other hand, was not significantly different than the Japanese Sapporo population; we suggest that an admixture event that occurred in the North of France with flies from U.S.A could explain the difference between the two French populations [24]. The difference between the two American populations seemed odd at first; however, the transcriptomic analysis revealed that a copper detoxification pathway specific to the Watsonville population (as discussed below) could be involved. Even though we should confirm these findings with a larger sampling, our results demonstrate the importance of considering different populations in such studies.

### Basal gene expression is different between invasive and native genotypes

We performed a transcriptomic analysis on genotypes from each of the three sampled locations in order to identify molecular processes underlying variation in the oxidative stress response. We first identified DE genes between the three genotypes in untreated conditions (control).

In *D. melanogaster*, genotypic differences accounted for 7.3% of DE genes showing micro-environment plasticity among as set of 16 DGRP (Drosophila Genetic Reference Panel) lines reared under carefully controlled standard conditions [46]. This result is in agreement with our results, in which almost 7% of the transcriptome was differentially expressed across genotypes. Most of these DE genes correspond to biological processes such as metabolism or protein synthesis and may possibly reflect genotype-specific differences related to local adaptation. In general, the level of expression for DE genes in invasive genotypes (U.S.A and France) was lower than in the native genotype from Japan, suggesting this genotype has by default a higher level of transcription for the DE genes. Among down-regulated genes, we found significant enriched GO terms in invasive genotypes related to translation, protein metabolic process, ribosome biogenesis, response to hyperoxia and immune response.

### Relationship between oxidative stress response, phenotype and gene expression

Exposure to paraquat affected the expression of up to 5% of the transcriptome (703 DE genes between control and paraquat) with a majority of DE genes being up-regulated. Similar changes in gene expression have been observed in *D. melanogaster*, with 608 to 1111 DE genes identified after exposure to 5mM or 15mM of paraquat [47]. In response to oxidative stress following exposure to hydrogen peroxide, 1639 DE genes were identified [47,48].

Interestingly, the proportion of the transcriptome affected by oxidative stress differed between native and invasive genotypes. The Japanese genotype appeared highly stable, with fewer DE genes in response to paraquat than both invasive genotypes (Fig. 4). Furthermore, the number of DE genes uniquely affected by paraquat exposure was much lower in the Japanese genotype (28) than either the U.S.A (96) or France (327) genotypes.

It should be pointed out that the two invasive lines used in the transcriptomic analysis, Watsonville (USA) and Montpellier (France), were the lines most affected by paraquat exposure in our phenotypic analysis (Fig. 1), suggesting that stress sensitivity could be linked to greater transcriptional deregulation. The French genotype had by far the greatest number of DE genes (almost twice as many as the American genotype). Consistent with the hypothesis of transcriptional deregulation, this result could reflect a lack of adaptation to paraquat, which has been banned as an herbicide in Europe since 2007, prior to the arrival of *D. suzukii*.

A set of 67 DE genes were shared by all genotypes in their response to paraquat. This set of common DE genes likely corresponded to those directly implicated in stress response. In agreement with this idea, we founded genes such as *Hsp* or genes of the cytochrome gene family [49–54]. At the transcriptomic level, the Japanese genotype appeared dissimilar from the other two. First, as discussed above, the transcriptional response to paraquat involved a much smaller portion of the genome and there were fewer DE genes unique to this genotype. Second, under control conditions, this genotype had the highest amount of DE genes. At the phenotypic level, the Japanese genotype had the lowest lifespan under standardized control conditions but was one of the genotypes most resistant to oxidative stress.

Together, these results suggested that the Japanese genotype maintained some constitutive defences to oxidative stress. In the absence of oxidative stress, the expression of constitutive defence may come at a cost of reduced lifespan but it would result in greater resistance when flies encounter paraquat. In the case of the French and American genotypes, many up-regulated genes are directly related to the oxidative stress response (GO enriched for oxidation-reduction, immune response and ion binding), which could indicate they are experiencing a greater amount of oxidative damage and therefore explain their lower lifespan under stress conditions.

A surprising result comes from the GO analysis of DE genes in response to paraquat exposure in the U.S.A genotype. We identified an enrichment in terms related to copper detoxification that was not found in the other genotypes. Previous studies, ranging from bacteria to mammals, have demonstrated a trade-off between copper tolerance and sensitivity to paraquat [55,56]. Thus, the greater sensitivity of the Watsonville genotype to paraquat exposure could reflect previous exposure and adaptation to copper in the environment. In support of this hypothesis, information from the California Pesticide Information Portal (https://calpip.cdpr.ca.gov/main.cfm) indicates a sizeable use of copper-based agricultural products, especially fungicides, with ∼525 Kg reportedly used in 2017.

### Genotype-specific transcriptional responses to paraquat exposure

To better understand genotype-specific responses to paraquat exposure, we focused on genes presenting a genotype-by-environment interaction (GEI). If differences in the transcriptional response to paraquat exposure reflect adaptive changes involved in response to local environmental conditions then analysis of genes with GEI may provide insight into the mechanisms of local adaptation [57]. Genes with GEI have often been identified in studies of oxidative stress responses (see *e*.*g*., Jordan *et al*., 2012) [58].

Genetic variation in transcriptomic plasticity could contribute to rapid adaptation to novel environments during the invasive process, possibly due to variation in both *cis* and *trans* regulatory sequences [52,59–63]. We founded evidence of GEI for the transcriptional response to paraquat in only a small part of the transcriptome. Most DE genes with evidence of GEI showed a greater change in the level of expression in invasive genotypes versus the native one. Due to the large number of genes that remain unannotated in the *D. suzukii* genome, a complete scenario of the genome-wide transcriptional response to oxidative stress is difficult to achieve. This may be particularly problematic when attempting to understand the functional relevance of genotype-specific responses. However, our results confirmed that for parts of the genome, the transcriptional response to oxidative stress varies across genotypes, and that some of these differences may reflect population history. Interestingly, the French genotype showed massive up-regulation of some *Hsp*’s compared to the USA and Japan genotypes (Fig. 7). These genes are known to be highly responsive to temperature [52] and also to oxidative stress (see review [49]). In general, GO analysis revealed an enrichment in terms related to oxidative stress (oxygen level, hyperoxia, hypoxia, stress) for up-regulated genes in invasive genotypes relative to the native one. These GO terms were also enriched for up-regulated genes with evidence of GEI in the comparison between France and the U.S.A.

One caveat of the genome-wide expression analysis is our statistical power to identify biologically relevant differences in expression levels. We have applied a threshold (FDR < 0.01 and absolute log_2_FC > 1) to identify DE genes, but it is possible that genes with more subtle changes in expression are important. Indeed, genes showing evidence of GEI are often found in upstream parts of regulatory networks, where even very small differences in expression could have pronounced phenotypic consequences. Also, genes with GEI are often associated with genetic variation in cis or trans-regulatory sequences, and a further investigation would be necessary to identify such factors in our data [52,62–65].

### TE insertions are depleted near oxidative stress sensitive genes

TEs have been described as stress sensitive and their activation by stress-responsive elements (SREs) in promoting regions could generate a burst of transposition and facilitate adaptation by increasing the genetic diversity upon which selection could act [13,66]. A recent review cited several examples of TE family activation following stress, which may depend on the type of stress and the TE family [17,67]. Horváth *et al*. (2017) also suggested that under stressful conditions, some TEs could be repressed just after their activation, indicating that stress could induce both activation and repression. TE transcription is a prerequisite to TE activity [17]. Our analysis of the TE transcriptome after stress induction showed that in *D. suzukii* very few TEs are activated, with a maximum of 6 TE families deregulated following exposure to paraquat in the French genotype. This result is not related to the potential activity of TE in *D. suzukii*, since a greater number of TE families are DE between genotypes in control conditions, suggesting that TEs in *D. suzukii* are capable of being expressed.

Most TE insertions are neutral or slightly deleterious, but some may be beneficial and implicated in adaptation [13,14,66,67]. The impact of TE insertions is often achieved by their effect on gene expression, likely due to the addition of regulatory sequences, present in the TE, that can modulate genes expressed, particularly during stress [20]. While TEs have been revealed as playing an important role in the success of invasive species, by generating genetic diversity and thus compensating for bottleneck effects after introduction, no empirical data exists to support this hypothesis [3,6,16,66].

As ∼33% of the *D. suzukii* genome is composed of TEs, we tested the hypothesis that TEs could modulate gene expression by the addition of regulatory regions. We found that the distribution of insertions along the chromosome did not differ among the three genomes, and, as observed in other Drosophila species, a majority of the insertions were in intergenic and intronic regions [68]. However, when we specifically analyzed DE genes, we observed a depletion of TE insertions, suggesting that TE insertions in stress response genes may be eliminated by strong purifying selection. This paucity of TE insertions was also observed for DE genes that were shared by the three genotypes. A gene encoding a glutathion-s-transferase was the only one to display a shared TE insertion in its flanking region. Finally, we tested for the enrichment of TE insertions in genes presenting GEI. We detected more insertions in genes with GEI than in other DE genes, suggesting that this category of gene may be more permissive to TE insertions. Several insertions were found in 3’UTR and 5’UTRs and could have regulatory impacts on those genes. Further analyses are needed to understand the molecular mechanisms responsible for changes in gene expression for this category of genes.

## Conclusion

Our results showed a difference in paraquat resistance between native and invasive populations of *D. suzukii*, that is not homogeneous between sampling sites on the same country. The differences observed between the two French populations could be explained by differential admixture subsequent to colonization in these two regions of France. In the United States, possible local adaptation to copper in the environment in Watsonville, as revealed by the functional analysis, may explain the difference in resistance to paraquat. Further research is required to test these hypotheses and to better understand population differences in paraquat resistance. Our data also reveal that gene expression patterns first depend on the genotype, and on the stress condition to a lesser extent. Finally, we showed that contrary to expectations, oxidative stress does not induce significant activation of TEs and that DE genes under stress conditions are depleted of TE insertions in the three genotypes of *D. suzukii* studied. Our results highlight that it is important to focus on several genotypes in performing phenotypic or transcriptomic analysis, and that we should consider the neglected role of TEs in adaptive evolution. Also, phenotypic and molecular approaches should complement each other to better understand the evolution of biological traits.

## Materials and Methods

### Drosophila suzukii lines rearing conditions and phenotyping

*D. suzukii* genotypes were sampled in 2014 in the native area (Japan: Sapporo and Tokyo) and two invaded areas (U.S.A: Watsonville and Dayton and France: Montpellier and Paris) (Table S8). To establish isofemale lines, a single gravid female was placed in a culture vial, and the line maintained thereafter with a low larval density in vials containing modified “Dalton” medium (Table S9) in a controlled environment: 22.5°C ± 1°C, 70 % ± 5% RH (relative hygrometry) and a 16:8 (Light/Day) [69]. We used paraquat (methyl viologen dichloride hydrate, ref. 75365-73-0, Sigma-Aldrich^®^) to mimic oxidative stress. Paraquat (10mM) was added to the cooling medium, before pouring into vials. Control vials were made at the same time but without adding paraquat. In the experiment, ten 4-7-day old flies were placed in experimental vials and transferred to new vials every 3 to 4 days to limit microbial development. Both males and females were tested and kept in separate vials. Survival was monitored by visual inspection every 24h. There were three replicate vials for each combination of the 27 isofemale lines (Table S7), sex, and paraquat treatment, for a total of 324 vials.

### Survival data analysis

The analysis of survival data was performed in two steps on R software (v.3.6.0, [70]). First, for each replicate (10 survival times), we used the fitdistcens function from the fitdistrplus package (v.1.0-14, [71]) to determine which of several distribution models (Weibull, lognormal and gamma) were most appropriate to r fit our right censored data (33 flies) data. The Weibull distribution was chosen after graphical comparison with others, also confirmed using loglikelihoods of the fitted models. For each replicate the fitted distribution was summarized using its theoretical median. Second, a linear mixed model was fitted to the log transformed medians using the lmer function of the lme4 R package (v.1.1-21, [72]), and p-values were estimated using lmerTest (v.3.1-0, [73]) with treatment, sex and population (the 6 sampled cities) entered as fixed factors and isofemale line as a random factor). The main effect of sex and interactions with both treatment and population were removed after AIC comparison from the final model for analysis. The interaction between population and treatment (GEI effect) was kept in the model. Model coefficients are reported with their confidence intervals (0.95) in Table S9 and after exponential transformation on Fig. 1. These effects can be interpreted as multiplicative effect on the median lifespan compared to a reference, here chosen as the non-exposed group from Sapporo. So, for example, with the untreated Sapporo flies centered on 1, an effect of 0.2 for paraquat-treated Sapporo flies means they have 20% of the survival time of Sapporo flies without paraquat. Normality and homoscedasticity of residuals and normality of random effects were confirmed graphically after logarithmic transformation of median survival times. We also examined the correlation across the isofemale lines between log-transformed survival times in control and paraquat-treated conditions using a Pearson correlation coefficient in R (Fig. S1).

### DNA extraction and sequencing

We sequenced genomic DNA for one isofemale line per country: S29, W120 and MT47 respectively from Sapporo (Japan), Watsonville (U.S.A) and Montpellier (France). DNA was extracted using phenol chloroform from a pool of 10 adult females. Libraries and sequencing were performed by the platform GeT-PlaGe, Génopole Toulouse / Midi-pyrénées (France), using Illumina (150 bp) *TruSeq Nano pair end*. We obtained between 33,362,864 and 72,022,388 reads per library. Sequences were cleaned using Trimmomatic with default parameters [74].

### RNA extraction and sequencing

We used the same three isofemale lines (S29, W120 and MT47) for our analysis of gene expression. For each of two biological replicates, fifteen 4-7 days old females were exposed for 24h to medium supplemented with paraquat (20mM) or without paraquat (*i*.*e*., a total of 12 samples). Flies were dissected on ice in a phosphate buffer saline solution to remove gonads, and the remaining somatic tissue was frozen in liquid nitrogen and stored at −80°C.

We used the RNAeasy Plus Mini Kit (Qiagen) to extract total RNA from the somatic tissues, following the protocol provided by manufacturer. Samples were treated with DNAse (ref AM2224, AMbion™) according to manufacturer instructions and stored at −80°C. RNA amount and quality was estimated using Qubit™ (Thermo Fisher Scientific) and the 2100 Bioanalyser instrument (Agilent). RNA libraries and sequencing were performed on the GenomEast platform, a member of the ‘France Génomique’ consortium (ANR-10-INBS-0009). Libraries were constructed using the TruSeq® Stranded mRNA Library Prep Kit following manufacturer’s recommendations. The libraries were sequenced on Illumina High HiSeq 4000 with paired-end 100 base pair long reads.

### Transcriptome analysis

Between 62.76 to 120.12 million paired-end reads were generated from the 12 libraries. Quality was assessed using FastQC (v. 0.10.1), a trimming step implemented with UrQt (v. 1.0.17, minimum phred score of 20), and quality was checked again using FastQC [75,76]. RNA-seq data were mapped on the *D. suzukii* reference genome using HISAT2 (v. 2-2.1.0) and read counts for genes were computed with eXpress [77–79]. We performed a reciprocal BLASTN (2.2.26) between the *D. suzukii* genes and the *Drosophila melanogaster* database (FlyBase, dm6 version) (archive data: FB2018_06) in order to identify orthologues [80]. Another BLASTX was performed against the NCBI nr database, using predicted genes *in D. suzukii* for which no orthologues were detected in *D. melanogaster*. Matched hits from this BLASTX were tagged with the term “(predicted)”. Of the 16905 annotated genes in the *D. suzukii* genome, 8428 matched with a Flybase gene and 478 others on the nr database (52.7% of total genes).

Differential expression analysis was performed using the DESeq2 package (v. 1.24.0) on R (v. 3.6.0) [81]. We built a model estimating the effects of genotype (France, U.S.A and Japan), the environment (control and paraquat), and the genotype-by-environment interaction (GEI effect). The *lfcShrink* function was used to estimate log_2_fold change and identify differentially expressed (DE) genes using the ashr R package [82]. DE genes were those with an FDR-adjusted p-value below 0.01 and absolute log_2_fold change > 1. The coefficient of variation (CV, standard deviation/mean) on normalized counts was computed for each genotype, between control and paraquat.

### Transposable element (TE) identification

The reference genome was masked using a custom TE library (Mérel *et al*., *in prep*). The Python script create-reads-for-te-sequences.py was used to generate reads corresponding to the TE library using the following parameters : *—read-length 125, --max-error-rate 0, --boost 10*) [77,83]. The reads were then mapped to the reference genome using bwa bwasw (v0.7.17) [84]. Aligned bases were masked using bedtools, bamtobed, and bedtools maskfasta (v2.20.0) [85]. This process of read generation and mapping was repeated 200 times. Note that sequences smaller than 500 bp were removed from the TE library. Forward and reverse reads were mapped separately to a fasta file containing the masked reference genome and the TE library. The mapping was done using bwa bwasw. For each line, the resulting single-end read alignments files were merged using PoPoolationTE2 se2pe (v1.10.04) [83]. PoPoolationTE2 pipeline was used to estimate TE frequencies in each sample. The following options were used in the analysis: *--map-quality 15* (ppileup module), *--mode joint, -- signature-window minimumSampleMedian, min-valley minimumSampleMedian, --min-count 2* (identify signature module), *--max-otherte-count 2, --max-structvar-count 2* (filterSignatures module), *--min-distance −200, --max-distance 300* (pairupSignatures module). In the PoPoolationTE2, hierarchy file was a file allowing multiple slightly diverged sequences to be assigned to one family, and all sequences with cross mapping reads were regrouped in the same family. The cross mapping was investigated by generating TE reads using create-reads-for-te-sequences.py (*--read-length 125, --max-error-rate 0, -- boost 50*) and mapping the reads to the TE library using bwa bwasw.

The software was run using the S29, W120 and MT47 DNAseq data. Using the gene annotation of the reference *D. suzukii* genome we identified TE insertions present in genes (exon, intron, 5’ and 3’ UTR) and ± 2kb flanking regions.

We tested the dependence of TE insertions with the state of the genes (DE or not) using a Chi-square test. We considered as absent, TEs with insertion frequency < 0.2 and present when > 0.8. Intermediate frequencies were removed to limit bias. For studies of TE insertions and expression of DE genes, we considered a potential effect of an insertion when frequency > 0.5.

### TE expression analysis

TE expression was quantified using the TEcount module from the TEtools software [86]. Briefly, TEcount sums reads aligned against copies of each TE family annotated from the reference genome creating an output table of expression arranged by TE family [77]. Differential expression of TEs between paraquat-treated and control flies for each isofemale line was computed using a merged file with the RNA counts for genes and TE families, and following normalization using DEseq2.

### TFBS screening

TE sequences inserted in flanking regions located ± 2kb from differentially expressed genes were screened for transcription factor binding sites (TFBS). We selected three TFBS (CNC, HSF and DL) related to antioxidant response element family (ARE) from the literature [41]. TFBS were screened in R (v. 3.6.0) using the JASPAR2018 database R library (v.1.1.1) and TFBSTools R library (v.1.22.0) [87,88]. PFM (Position Frequency Matrix) matrices were extracted (CNC:MA0530.1, HSF:MA0486.2, DL:MA0022.1) before a PWM (Position Weight Matrix) conversion with the *pseudocount* value set to 0.8. The minimum score value for the screening was fixed at 0.95 to minimize false positives due to small TFBS sequence sizes. P-values were adjusted with the Benjamini-Hochberg correction for multi-testing [89].

### Gene ontology analysis

We performed a GO enrichment analysis directly on the geneontology.org website, using homologs in *D. melanogaster* to discover over or underrepresented gene functions from the lists of DE genes [54].

P-values were calculated using a Fisher test for enriched GO terms and adjusted with the Benjamini-Hochberg correction for multi-testing [89]. GO terms with FDR ≤ 0.05 were defined as significantly enriched. The GO terms were reduced to representative non-redundant terms using the REVIGO tool and manual curation [90].

## Declarations

## Acknowledgment

Experimental procedures were supported by the ANR (grant SWING ANR to P.G. and C.V. & grant ExHyb ANR to C.V.) and the Rovaltain foundation (EpiRip project). Bioinformatic work was performed using the computing facilities of the CC LBBE /PRABI. We thank S. Vicaire for the RNA libraries preparation, K. McKean for english editing and suggestions in the manuscript, N. Burlet, S. Martinez and H. Henri for various technical contributions.

## Contribution

P.M. produced data, conceived and wrote the manuscript draft. C.V. & P.G. designed experiments, edited the manuscript. J.P. trimmed NGS data and help in bioinformatic analysis. A.J. calibrated experimental design and produced data. M.L.D.M. supervised the statistical analysis and revised manuscript. M.F. helped on transcriptomic analysis and manuscript correction. M.G.F. performed gene ontology analysis. V.M. produced all related TE informations (genome annotations, frequency insertions). All authors proofread and approved the manuscript

## Ethics declarations

### Ethics approval and consent to participate

Not applicable.

### Consent for publication

Not applicable.

### Competing interests

The authors declare that they have no competing interests.

## Availability of data

DNA and RNA-seq data are available on NCBI with the Sequence Read Archive (SRA) number: SUB7319267 & SUB7021358.

Survival data of the populations are available to: ftp://pbil.univ-lyon1.fr/pub/datasets/Marin2020/surv_suzukii_marin2020/surv_suzukii_marin2020.txt

## Supplementary figures

**Fig. S1.**
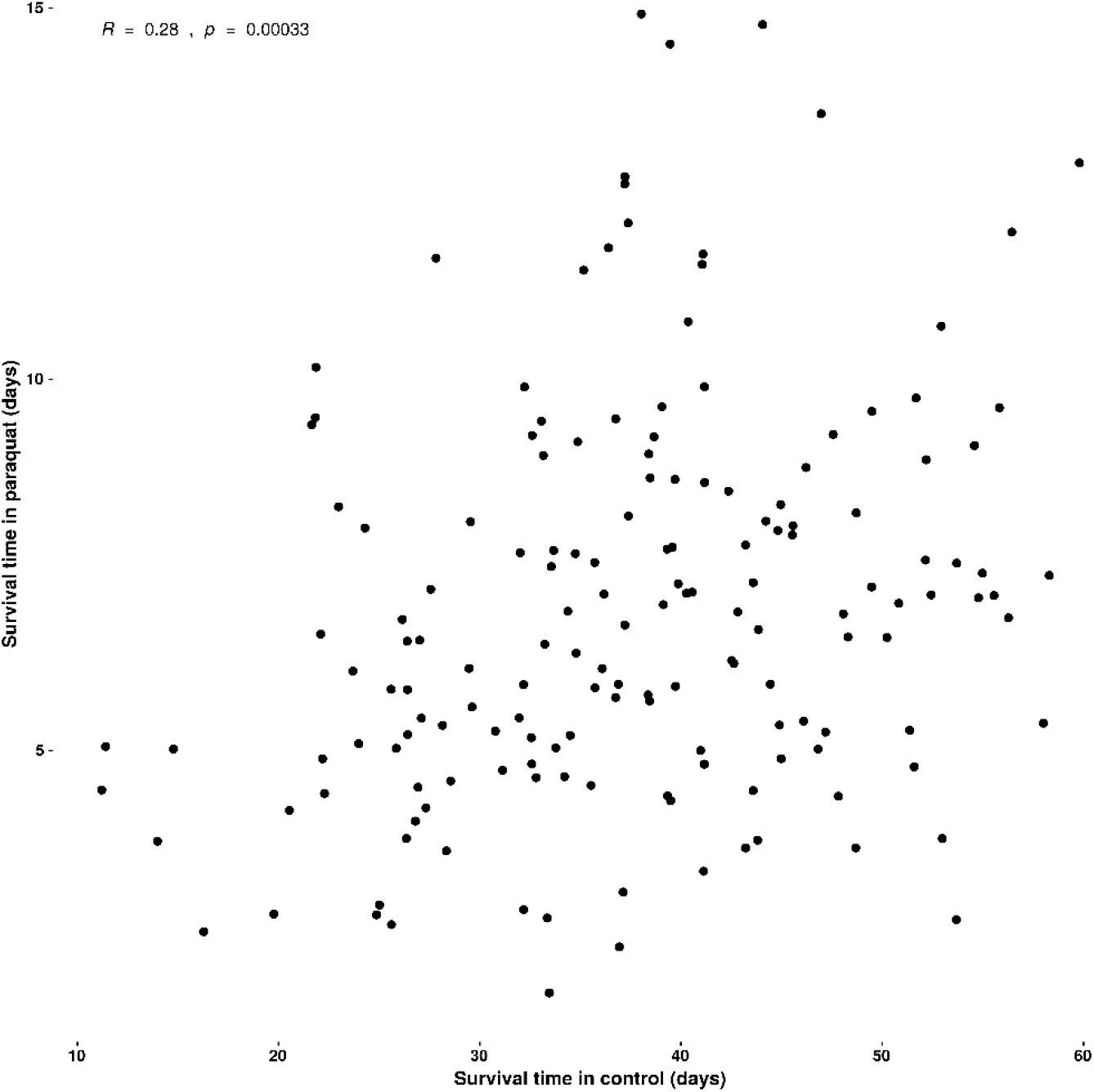
Correlation between median lifespan under or not paraquat treatment for every line. Correlation test was made using Pearson method.

**Fig. S2.**
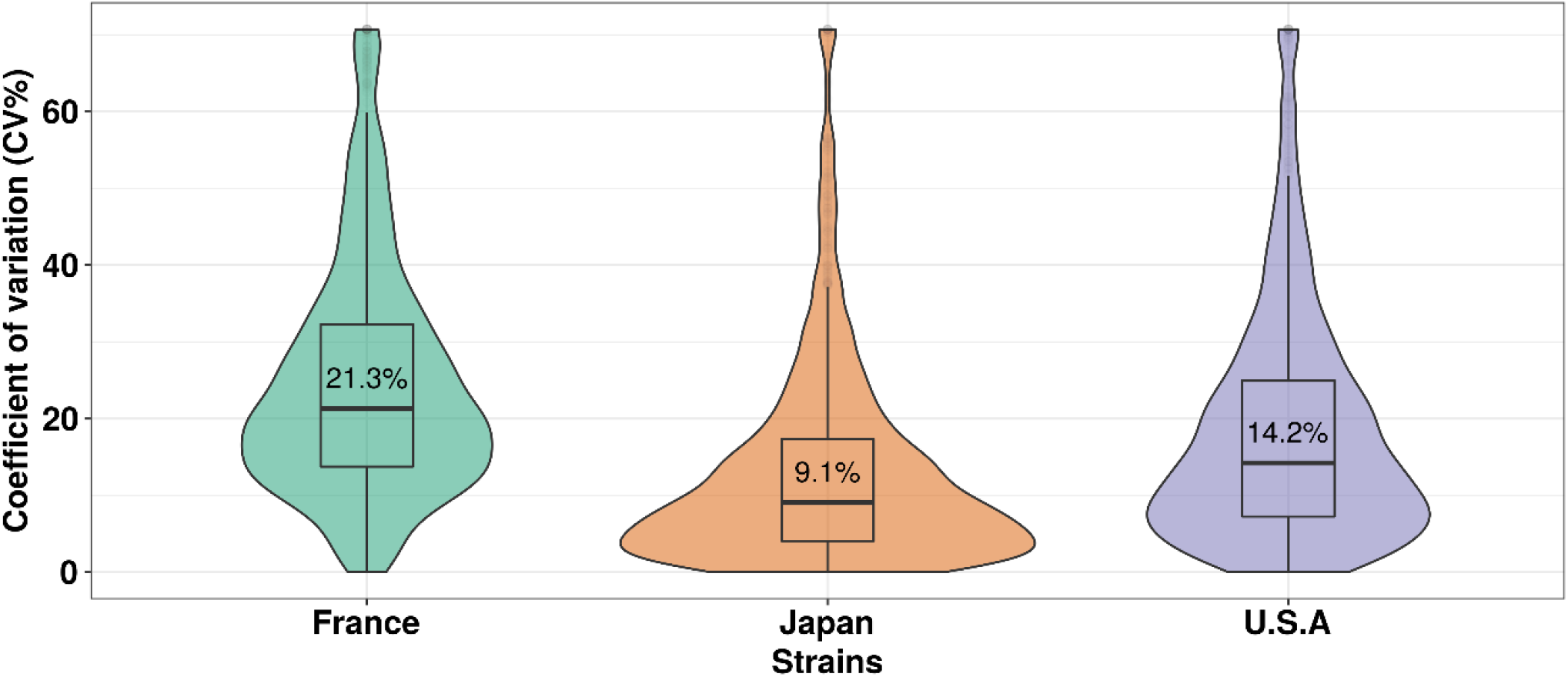
Distribution of the coefficient of variation (%) of DE genes after exposure to paraquat. Coefficients of variation was calculated using the standard deviation and mean counts in control and paraquat treated flies. Central values correspond to the median coefficient of variation. Pairwise comparisons of medians were done using a paired Wilcoxon test and all comparison were significants (p-value < 0.01).

**Fig. S3.**
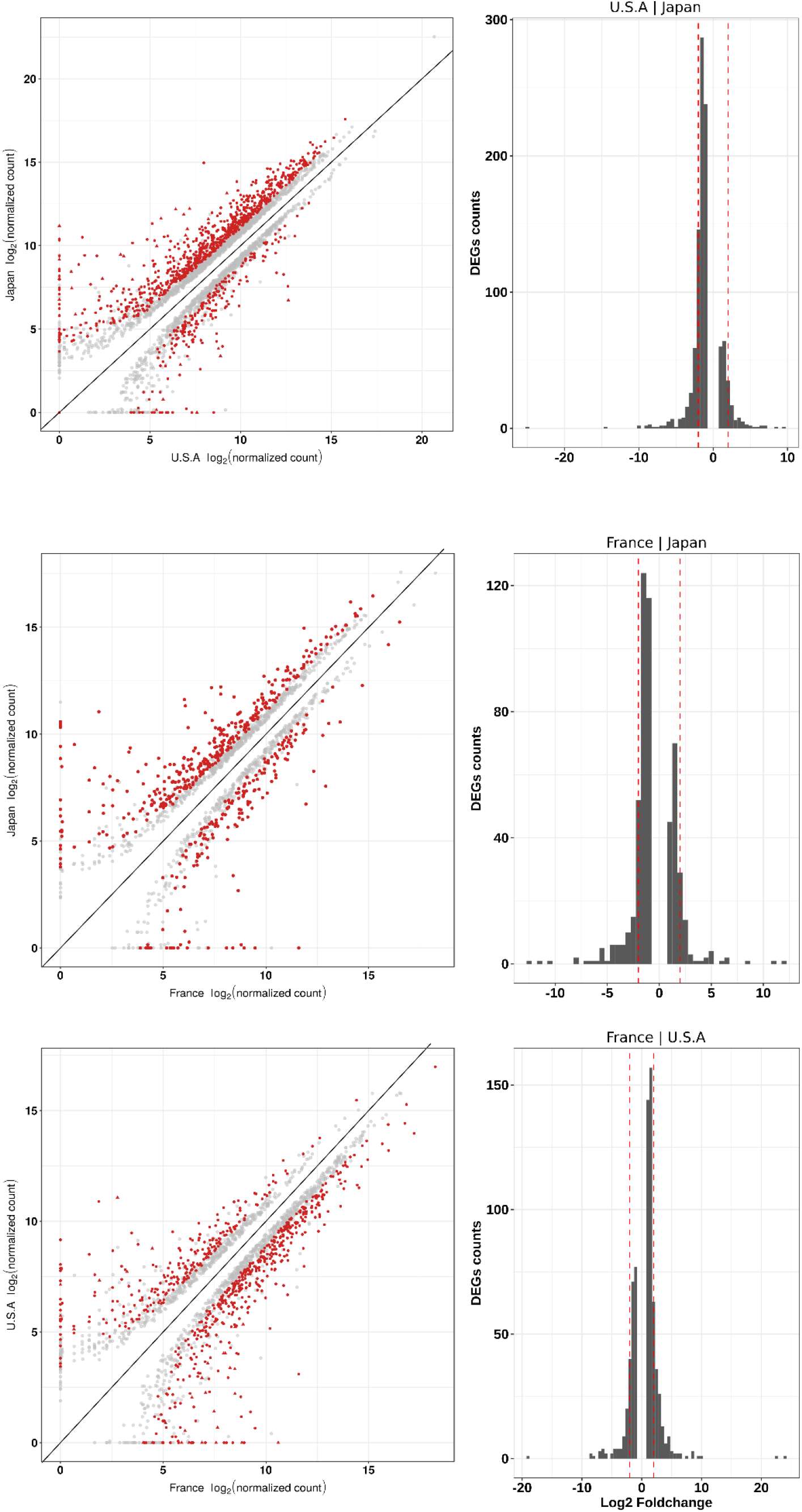
Scatter plots (left) of significant differentially expressed genes in pairwise comparisons between populations under control conditions using log_2_ of normalized counts. Histograms (right) of log_2_-fold-changes for DE genes in pairwise comparisons between populations under control conditions. Red lines correspond to threshold of fold change = 2.

**Fig. S4.**
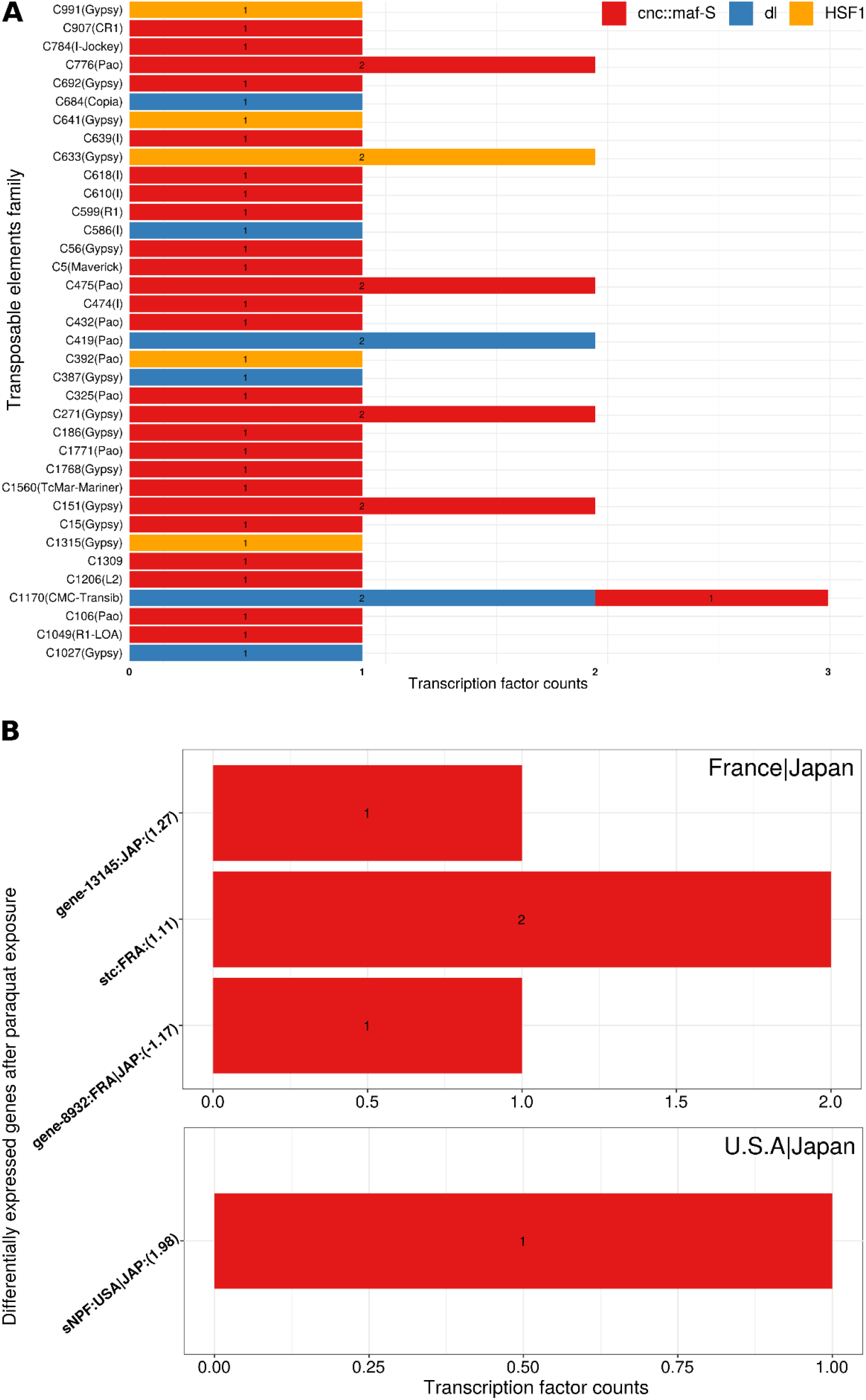
A Transcription factor binding site counts in candidate TEs. (B) DE genes identified from comparisons between populations after paraquat treatment (a GEI). Colours correspond to the 3 screened TFBS, CnC (red), dl (blue), and HSF1 (orange). Gene names are labelled on the y-axis, followed by the genotype where insertion is detected (FRA for France, JAP for Japan, USA for U.S.A and possible combination for shared), and the log_2_-fold-change of expression in contrasted genotypes.

## Supplementary tables

**Table S1.**
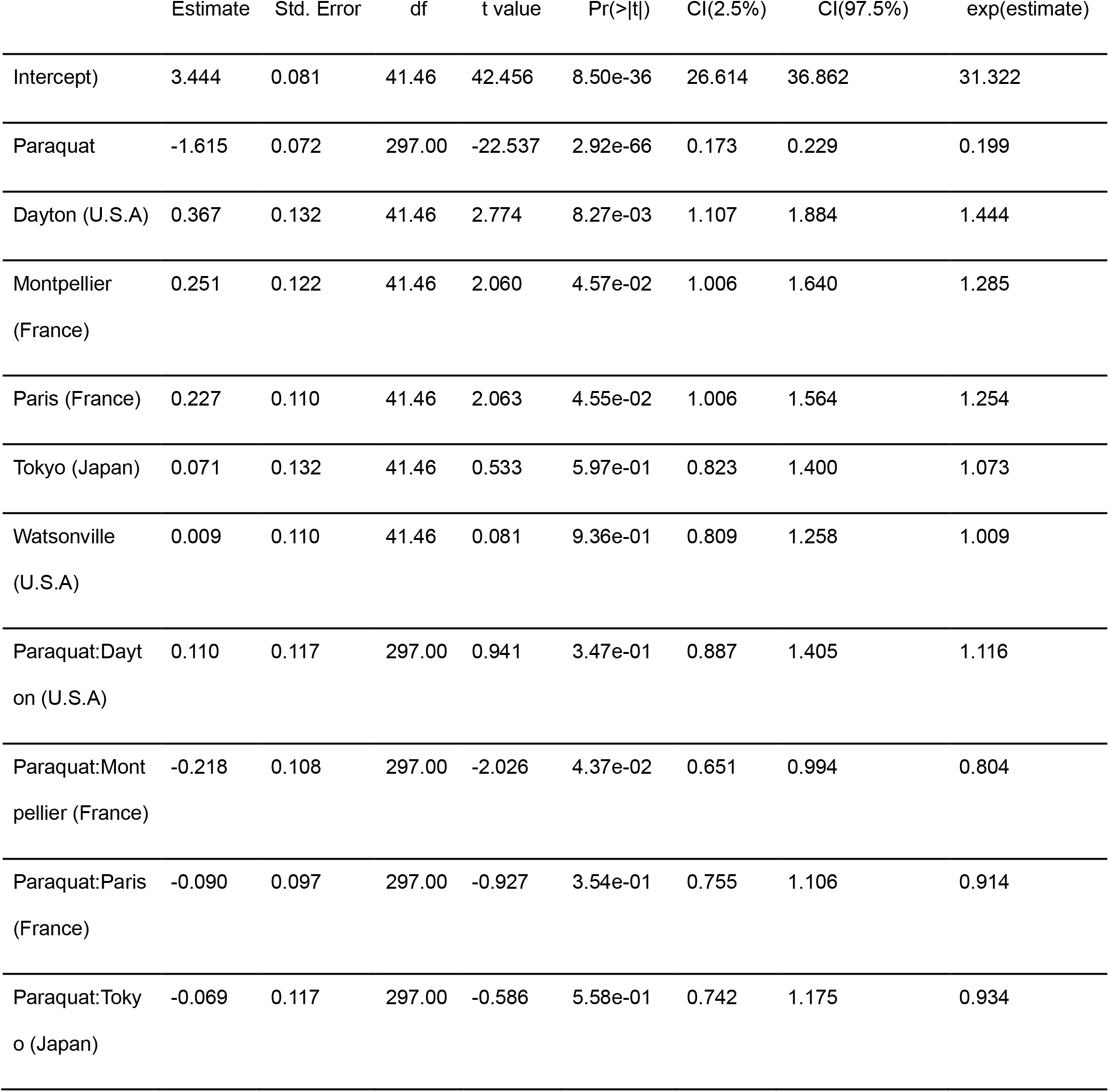
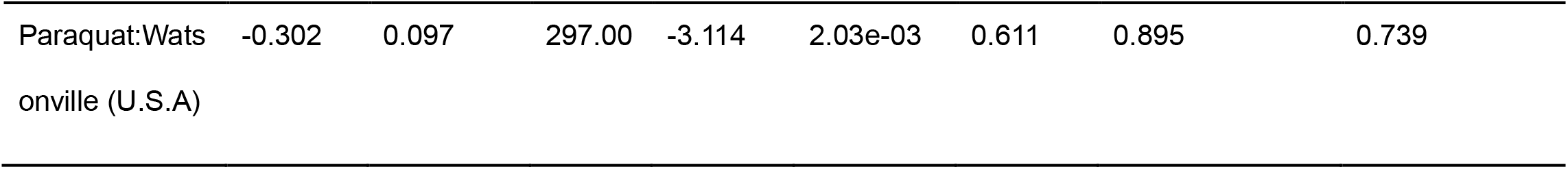
Table of linear mixed model with estimated coefficient and associated statistic. Model is centered on the Sapporo population reference, in untreated condition. Data were previously log transformed for normality. Isofemale lines were included as a random effect and we used exponential of coefficient value as multiplicative effect to interpret.

**Table S2.**
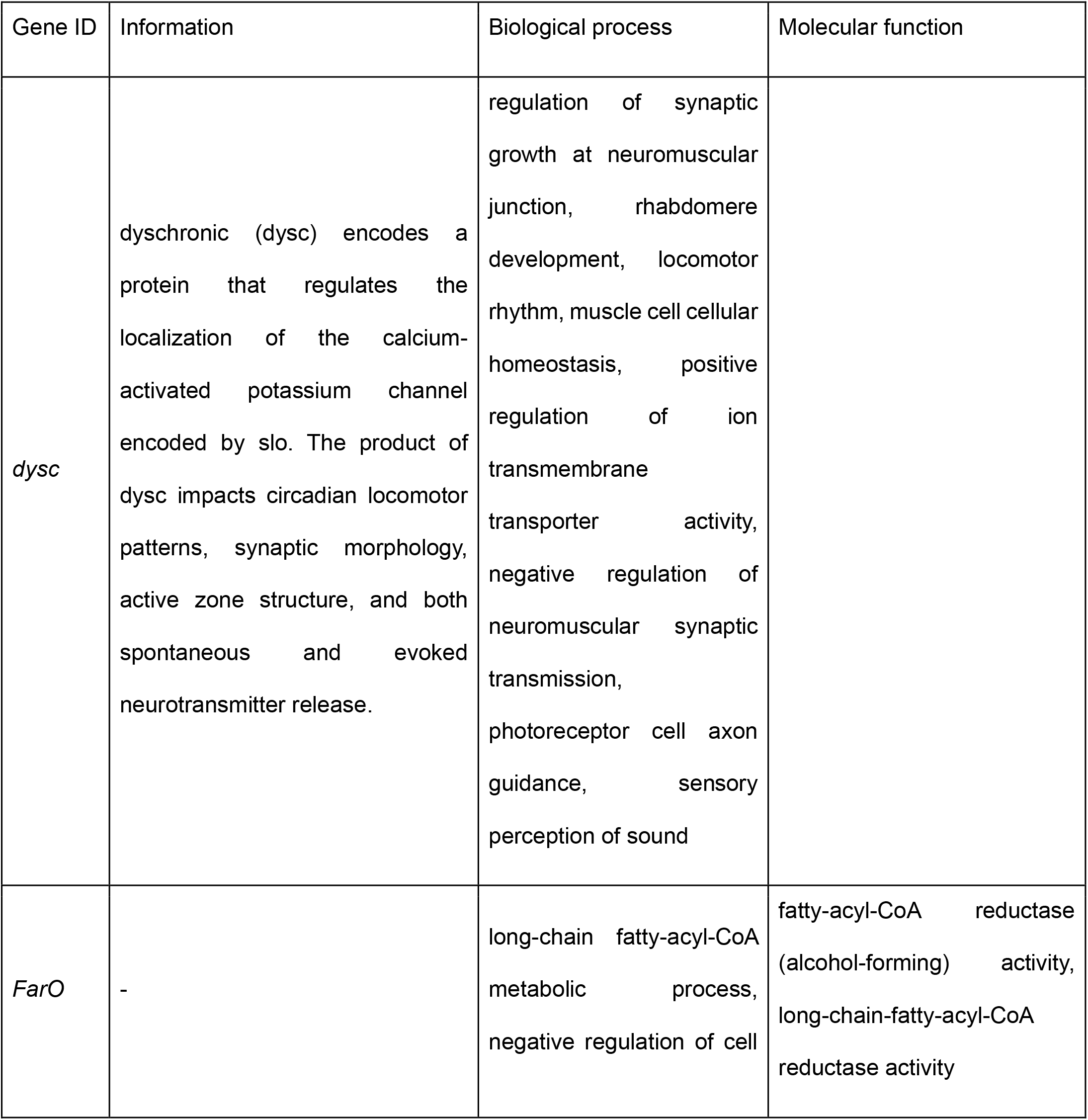
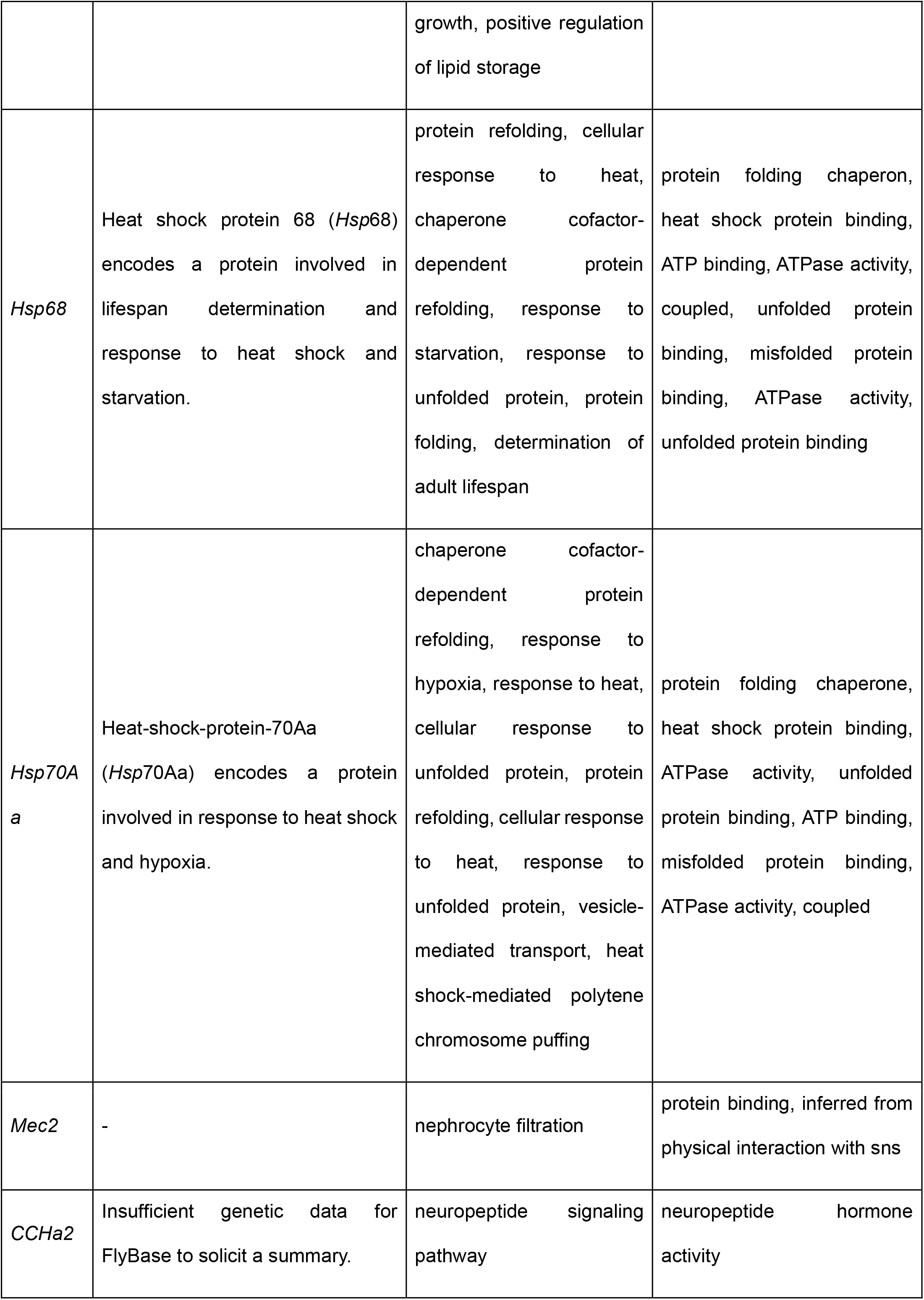
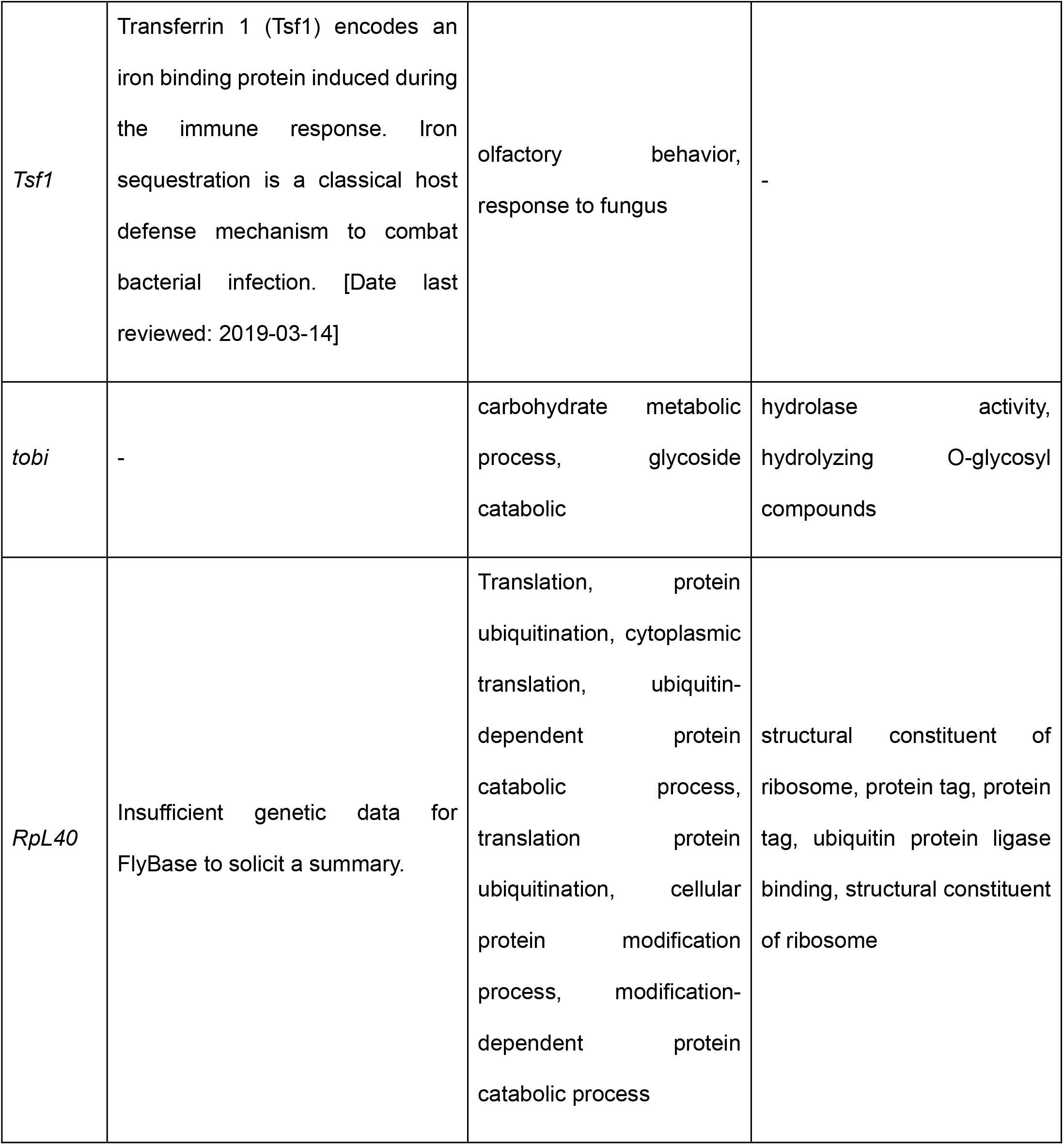
Table of some differentially expressed genes (p-adjusted <0.01 and log_2_FC > 1). These genes are exampled in Fig. 7. in the genotype environment interaction (GEI) with flybase information available.

| Gene ID | Information | Biological process | Molecular function |
| --- | --- | --- | --- |
| *dysc* | dyschronic (*dysc*) encodes a protein that regulates the localization of the calcium-activated potassium channel encoded by *slo*. The product of *dysc* impacts circadian locomotor patterns, synaptic morphology, active zone structure, and both spontaneous and evoked neurotransmitter release. | regulation of synaptic growth at neuromuscular junction, rhabdomere development, locomotor rhythm, muscle cell cellular homeostasis, positive regulation of ion transmembrane transporter activity, negative regulation of neuromuscular synaptic transmission, photoreceptor cell axon guidance, sensory perception of sound |  |
| *FarO* | - | long-chain fatty-acyl-CoA metabolic process, negative regulation of cell growth, positive regulation of lipid storage | fatty-acyl-CoA reductase (alcohol-forming) activity, long-chain-fatty-acyl-CoA reductase activity |
| *Hsp68* | Heat shock protein 68 (*Hsp68*) encodes a protein involved in lifespan determination and response to heat shock and starvation. | protein refolding, cellular response to heat, chaperone cofactor-dependent protein refolding, response to starvation, response to unfolded protein, protein folding, determination of adult lifespan | protein folding chaperon, heat shock protein binding, ATP binding, ATPase activity, coupled, unfolded protein binding, misfolded protein binding, ATPase activity, unfolded protein binding |
| *Hsp70Aa* | Heat-shock-protein-70Aa (*Hsp70Aa*) encodes a protein involved in response to heat shock and hypoxia. | chaperone cofactor-dependent protein refolding, response to hypoxia, response to heat, cellular response to unfolded protein, protein refolding, cellular response to heat, response to unfolded protein, vesicle-mediated transport, heat shock-mediated polytene chromosome puffing | protein folding chaperone, heat shock protein binding, ATPase activity, unfolded protein binding, ATP binding, misfolded protein binding, ATPase activity, coupled |
| *Mec2* | - | nephrocyte filtration | protein binding, inferred from physical interaction with sns |
| *CCHa2* | Insufficient genetic data for FlyBase to solicit a summary. | neuropeptide signaling pathway | neuropeptide hormone activity |
| *Tsf1* | Transferrin 1 (Tsf1) encodes an iron binding protein induced during the immune response. Iron sequestration is a classical host defense mechanism to combat bacterial infection. [Date last reviewed: 2019-03-14] | olfactory behavior,<br>response to fungus | - |
| *tobi* | - | carbohydrate metabolic<br>process, glycoside<br>catabolic | hydrolase activity,<br>hydrolyzing O-glycosyl<br>compounds |
| *RpL40* | Insufficient genetic data for FlyBase to solicit a summary. | Translation, protein<br>ubiquitination, cytoplasmic<br>translation, ubiquitin-<br>dependent protein<br>catabolic process,<br>translation protein<br>ubiquitination, cellular<br>protein modification<br>process, modification-<br>dependent protein<br>catabolic process | structural constituent of<br>ribosome, protein tag, protein<br>tag, ubiquitin protein ligase<br>binding, structural constituent<br>of ribosome |

**Table S3.** Number of DE TEs between control and oxidative (paraquat) condition for each genotype and between the different genotypes for both conditions. DE TE threshold made with adjusted p-value ≤0.01 and absolute log_2_foldchange ≥ 1. The rate corresponds to number of DE TE on total TE families (2030).

| Carcasses | DE TEs | Up-regulated | Down-regulated | DE rate (%) |
| --- | --- | --- | --- | --- |
| France\|Japan control | 48 | 22 | 26 | 3.08 |
| France\|U.S.A control | 78 | 10 | 68 | 5.01 |
| U.S.A\|Japan control | 92 | 70 | 22 | 5.91 |
| Japan (paraquat\|control) | 3 | 3 | 0 | 0.19 |
| France (paraquat\|control) | 6 | 6 | 0 | 0.39 |
| U.S.A (paraquat\|control) | 5 | 3 | 2 | 0.32 |
| France\|Japan paraquat | 1 | 1 | 0 | 0.06 |
| France\|U.S.A paraquat | 2 | 2 | 0 | 0.13 |
| U.S.A\|Japan paraquat | 0 | 0 | 0 | 0.00 |

**Table S4.** Observed genomic distribution of TE insertions in Japan, U.S.A, France.

|  | intergenic | +/-2kb<br>flanking | 5'UTR | 3'UTR | intron | exon |
| --- | --- | --- | --- | --- | --- | --- |
| France | 17142 | 2179 | 66 | 76 | 2469 | 115 |
| U.S.A | 19210 | 2399 | 69 | 78 | 2582 | 124 |
| Japan | 18924 | 2354 | 73 | 87 | 2687 | 133 |

**Table S5.** Contingency table (observed and expected) of DE genes and TE insertions detected toward 2kb for the three genotypes. P-value associated correspond to the Pearson chi-square test result. The three first rows correspond to DE genes in every genotypes after paraquat exposure and last 3 rows to GEI genes in every contrasted genotypes. Partial chi-square are in brackets.

|  |  | Observed |  | Expected |  |  |
| --- | --- | --- | --- | --- | --- | --- |
|  |  | TE- | TE+ | TE- | TE+ | p-value |
| France | DE- | 11501 | 2506 | 11528.0 | 2479.0 | 0.00216 |
|  | DE+ | 464 | 67 | 437.0 | 94.0 |  |
| Japan | DE- | 11664 | 2752 | 11675.2 | 2740.8 | 0.01321 |
|  | DE+ | 110 | 12 | 98.8 | 23.2 |  |
| U.S.A | DE- | 11678 | 2579 | 11692.5 | 2564.5 | 0.02761 |
|  | DE+ | 245 | 36 | 230.5 | 50.5 |  |
| France \| Japan | DE- | 11358 | 3042 | 11352.2 | 3047.8 | 0.2678 |
|  | DE+ | 103 | 35 | 108.8 | 29.2 |  |
| France \| U.S.A | DE- | 11491 | 2982 | 11499.4 | 2973.6 | 0.01566 |
|  | DE+ | 60 | 5 | 51.6 | 13.4 |  |
| U.S.A \| Japan | DE- | 11383 | 3093 | 11385.2 | 3090.8 | 0.5894 |
|  | DE+ | 51 | 11 | 48.8 | 13.2 |  |

**Table S6.** Insertion position of TEs in genes differentially expressed

|  |  | 5'UTR | Exon | Intron | 3'UTR | Flank 2kb |
| --- | --- | --- | --- | --- | --- | --- |
| France | Down | 0 | 1 | 17 | 0 | 8 |
|  | Up | 0 | 2 | 18 | 2 | 19 |
| U.S.A | Down | 0 | 0 | 6 | 0 | 5 |
|  | Up | 1 | 0 | 12 | 2 | 10 |
| Japan | Down | 0 | 0 | 2 | 0 | 3 |
|  | Up | 0 | 0 | 2 | 0 | 5 |

**Table S7.** GEI DE genes with inserted element, unknown gene in Flybase was reported with name “gene” followed by a number. Contrast column correspond to the lines tested for GEI. Type is the structure where is inserted the element with the position of right and inserted line correspond to the line where is detected the element.

| Gene symbol | log<sub>2</sub>FoldChange | Contrast | Type | Insertion position | Inserted line |
| --- | --- | --- | --- | --- | --- |
| *CG10383* | 1.753125 | France \| Japan | intron | 11431663 | Japan |
| *CG12520* | 1.892906 | France \| Japan | 3'UTR | 16312823 | Japan |
| *Mrp4* | 1.354537 | France \| Japan | intron | 3059393 | Japan |
| *FarO* | -4.923014 | France \| Japan | intron | 7509567 | France |
| *Ire1* | 1.168464 | France \| Japan | exon.part | 1288004 | Japan |
| *CG11626* | 1.328294 | France \| Japan | flank2kb | 1760496 | Japan |
| *(predicted)Hsp70-3* | 3.007337 | France \| Japan | intron | 2816351 | Japan |
| *Drgx* | 1.027039 | France \| Japan | intron | 23636644 | Japan |
| *Drgx* | 1.027039 | France \| Japan | flank2kb | 23660595 | U.S.A |
| *mesh* | 1.497837 | France \| Japan | intron | 1007389 | France |
| *Myo28B1* | 1.175073 | France \| Japan | 5'UTR | 7819926 | Japan |
| *Mocs1* | 1.115113 | France \| Japan | intron | 18050475 | U.S.A |
| *(predicted)kelch-like* | 2.399565 | France \| Japan | flank2kb | 14580423 | France |
| *stc* | 1.106598 | France \| Japan | flank2kb | 23035304 | France |
| *CG42808* | 2.321616 | France \| Japan | intron | 6914954 | France |
| *Pvf1* | 1.262606 | France \| Japan | flank2kb | 924620 | Japan |
| *Pvf1* | 1.262606 | France \| Japan | 5'UTR | 928977 | Japan |
| *Pvf1* | 1.262606 | France \| Japan | flank2kb | 924620 | France |
| *Pvf1* | 1.262606 | France \| Japan | 5'UTR | 928977 | France |
| *Pvf1* | 1.262606 | France \| Japan | flank2kb | 924620 | U.S.A |
| *Pvf1* | 1.262606 | France \| Japan | 5'UTR | 928977 | U.S.A |
| *JMJD4* | 1.152672 | France \| Japan | 5'UTR | 12034085 | U.S.A |
| *Ldh* | 1.414352 | France \| Japan | flank2kb | 2962505 | France |
| *Ldh* | 1.414352 | France \| Japan | flank2kb | 2962902 | France |
| *gene-8816* | -6.929561 | France \| Japan | exon.part | 826521 | Japan |
| *gene-8816* | -6.929561 | France \| Japan | flank2kb | 828928 | Japan |
| *gene-8816* | -6.929561 | France \| Japan | exon.part | 826521 | France |
| *gene-8816* | -6.929561 | France \| Japan | flank2kb | 828928 | France |
| *gene-8816* | -6.929561 | France \| Japan | flank2kb | 828928 | U.S.A |
| *CG4456* | 1.689628 | France \| Japan | flank2kb | 7586469 | U.S.A |
| *Oatp33Eb* | 1.420563 | France \| Japan | flank2kb | 2565015 | Japan |
| *Oatp33Eb* | 1.420563 | France \| Japan | flank2kb | 2565015 | France |
| *Oatp33Eb* | 1.420563 | France \| Japan | flank2kb | 2565015 | U.S.A |
| *Oatp33Eb* | 1.420563 | France \| Japan | flank2kb | 2566617 | U.S.A |
| *gene-16041* | -1.556090 | France \| Japan | intron | 51130 | Japan |
| *gene-16041* | -1.556090 | France \| Japan | intron | 53804 | Japan |
| *gene-16041* | -1.556090 | France \| Japan | intron | 51130 | France |
| *gene-16041* | -1.556090 | France \| Japan | intron | 53804 | France |
| *gene-16041* | -1.556090 | France \| Japan | intron | 51130 | U.S.A |
| *gene-16041* | -1.556090 | France \| Japan | intron | 53804 | U.S.A |
| *Hsp70Aa* | 1.956436 | France \| Japan | flank2kb | 2824573 | Japan |
| *Hsp70Aa* | 1.956436 | France \| Japan | flank2kb | 2824573 | U.S.A |
| *gene-742* | 1.419717 | France \| Japan | flank2kb | 2573404 | U.S.A |
| *CG3513* | 2.132744 | France \| Japan | flank2kb | 23660595 | U.S.A |
| *Socs36E* | 1.322709 | France \| Japan | flank2kb | 12039658 | France |
| *Socs36E* | 1.322709 | France \| Japan | intron | 12042814 | U.S.A |
| *Socs36E* | 1.322709 | France \| Japan | intron | 12046161 | U.S.A |
| *Socs36E* | 1.322709 | France \| Japan | intron | 12046748 | U.S.A |
| *gene-8522* | 1.051557 | France \| Japan | exon.part | 8449067 | Japan |
| *gene-8522* | 1.051557 | France \| Japan | exon.part | 8449067 | France |
| *gene-8522* | 1.051557 | France \| Japan | exon.part | 8449067 | U.S.A |
| *Drat* | 1.079845 | France \| JapanF | intron | 2761492 | France |
| *gene-12098* | 1.386302 | France \| Japan | intron | 6814225 | Japan |
| *lncRNA:CR45936* | -1.419969 | France \| Japan | intron | 16923423 | U.S.A |
| *gene-2030* | 1.575219 | France \| Japan | intron | 5579171 | Japan |
| *gene-2030* | 1.575219 | France \| Japan | intron | 5579171 | France |
| *gene-2030* | 1.575219 | France \| Japan | intron | 5579171 | U.S.A |
| *gene-3647* | 1.941648 | France \| Japan | intron | 2104897 | France |
| *gene-3647* | 1.941648 | France \| Japan | intron | 2099877 | U.S.A |
| *gene-1513* | 1.070880 | France \| Japan | intron | 145053 | France |
| *(predicted)trypsin_alpha3* | -2.153081 | France \| Japan | intron | 6743283 | France |
| *gene-13286* | 1.164912 | France \| Japan | 3'UTR | 16312823 | Japan |
| *gene-13145* | 1.267960 | France \| Japan | intron | 16350711 | Japan |
| *gene-13145* | 1.267960 | France \| Japan | intron | 16351595 | France |
| *Chs2* | 1.075446 | France \| Japan | flank2kb | 22441443 | Japan |
| *Chs2* | 1.075446 | France \| Japan | flank2kb | 22441443 | U.S.A |
| *gene-12969* | 1.158750 | France \| Japan | flank2kb | 18466778 | Japan |
| *gene-8932* | -1.170218 | France \| Japan | intron | 5844 | Japan |
| *gene-8932* | -1.170218 | France \| Japan | intron | 5844 | France |
| *gene-8932* | -1.170218 | France \| Japan | intron | 5844 | U.S.A |
| *gene-5432* | -1.652789 | France \| Japan | intron | 9876320 | France |
| *betaTub60D* | 1.037268 | France \| Japan | 3'UTR | 12121904 | France |
| *gene-16855* | -1.081314 | France \| Japan | flank2kb | 439861 | Japan |
| *gene-16855* | -1.081314 | France \| Japan | flank2kb | 440304 | Japan |
| *gene-16855* | -1.081314 | France \| Japan | flank2kb | 439861 | France |
| *gene-16855* | -1.081314 | France \| Japan | flank2kb | 440304 | France |
| *gene-16855* | -1.081314 | France \| Japan | flank2kb | 439861 | U.S.A |
| *gene-16855* | -1.081314 | France \| Japan | flank2kb | 440304 | U.S.A |
| *gene-16070* | -1.190533 | France \| Japan | intron | 45349 | Japan |
| *gene-16070* | -1.190533 | France \| Japan | intron | 53781 | Japan |
| *gene-16070* | -1.190533 | France \| Japan | intron | 56597 | Japan |
| *gene-16070* | -1.190533 | France \| Japan | intron | 92529 | Japan |
| *gene-16070* | -1.190533 | France \| Japan | intron | 97953 | Japan |
| *gene-16070* | -1.190533 | France \| Japan | intron | 99403 | Japan |
| *gene-16070* | -1.190533 | France \| Japan | intron | 118960 | Japan |
| *gene-16070* | -1.190533 | France \| Japan | intron | 45349 | France |
| *gene-16070* | -1.190533 | France \| Japan | intron | 53781 | France |
| *gene-16070* | -1.190533 | France \| Japan | intron | 56597 | France |
| *gene-16070* | -1.190533 | France \| Japan | intron | 92529 | France |
| *gene-16070* | -1.190533 | France \| Japan | intron | 97953 | France |
| *gene-16070* | -1.190533 | France \| Japan | intron | 99403 | France |
| *gene-16070* | -1.190533 | France \| Japan | intron | 118960 | France |
| *gene-16070* | -1.190533 | France \| Japan | intron | 45349 | U.S.A |
| *gene-16070* | -1.190533 | France \| Japan | intron | 53781 | U.S.A |
| *gene-16070* | -1.190533 | France \| Japan | intron | 56597 | U.S.A |
| *gene-16070* | -1.190533 | France \| Japan | intron | 92529 | U.S.A |
| *gene-16070* | -1.190533 | France \| Japan | intron | 97953 | U.S.A |
| *gene-16070* | -1.190533 | France \| Japan | intron | 99403 | U.S.A |
| *gene-16070* | -1.190533 | France \| Japan | intron | 118960 | U.S.A |
| *CG33282* | 1.002304 | France \| Japan | intron | 19704856 | U.S.A |
| *CG10361* | 1.045731 | France \| Japan | flank2kb | 18922002 | Japan |
| *(predicted)Hsp70-3* | 4.281151 | France \| U.S.A | intron | 2816351 | Japan |
| *Hsp70Aa* | 4.230943 | France \| U.S.A | flank2kb | 2824573 | Japan |
| *Hsp70Aa* | 4.230943 | France \| U.S.A | flank2kb | 2824573 | U.S.A |
| *FarO* | -5.293578 | France \| U.S.A | intron | 7509567 | France |
| *(predicted)kelch-like* | 2.863579 | France \| U.S.A | flank2kb | 14580423 | France |
| *CG10621* | -1.446417 | France \| U.S.A | 3'UTR | 11134738 | U.S.A |
| *gene-5842* | -1.775330 | France \| U.S.A | intron | 9902605 | Japan |
| *gene-5842* | -1.775330 | France \| U.S.A | intron | 9902872 | Japan |
| *gene-5842* | -1.775330 | France \| U.S.A | intron | 9905803 | France |
| *CG16965* | -1.049352 | France \| U.S.A | flank2kb | 3227015 | Japan |
| *gene-9109* | 6.739623 | France \| U.S.A | intron | 2816351 | Japan |
| *Ir48b* | 2.818729 | France \| U.S.A | intron | 10573980 | Japan |
| *Ir48b* | 2.818729 | France \| U.S.A | intron | 10573980 | U.S.A |
| *Tsf1* | -1.099044 | France \| U.S.A | intron | 1957700 | Japan |
| *Ldh* | 2.339272 | U.S.A \| Japan | flank2kb | 2962505 | France |
| *Ldh* | 2.339272 | U.S.A \| Japan | flank2kb | 2962902 | France |
| *CG12520* | 1.480822 | U.S.A \| Japan | 3'UTR | 16312823 | Japan |
| *sNPF* | 1.983076 | U.S.A \| Japan | intron | 14521385 | Japan |
| *sNPF* | 1.983076 | U.S.A \| Japan | intron | 14523939 | Japan |
| *sNPF* | 1.983076 | U.S.A \| Japan | intron | 14521385 | France |
| *sNPF* | 1.983076 | U.S.A \| Japan | intron | 14521385 | U.S.A |
| *CG4456* | 2.260613 | U.S.A \| Japan | flank2kb | 7586469 | U.S.A |
| *Mal-A1* | 2.109920 | U.S.A \| Japan | flank2kb | 5265200 | France |
| *CG16965* | 1.430053 | U.S.A \| Japan | flank2kb | 3227015 | Japan |
| *Mal-A7* | 2.224882 | U.S.A \| Japan | intron | 5210868 | Japan |
| *Mal-A7* | 2.224882 | U.S.A \| Japan | intron | 5211873 | Japan |
| *Mal-A7* | 2.224882 | U.S.A \| Japan | intron | 5211873 | France |
| *Mal-A7* | 2.224882 | U.S.A \| Japan | intron | 5211873 | U.S.A |
| *gene-3840* | 1.106626 | U.S.A \| Japan | intron | 1916531 | U.S.A |
| *tup* | 1.030389 | U.S.A \| Japan | intron | 11241399 | Japan |
| *DptB* | 2.623785 | U.S.A \| Japan | intron | 4000182 | U.S.A |
| *CG4372* | 1.269428 | U.S.A \| Japan | flank2kb | 1467799 | France |
| *Ance-2* | 1.004916 | U.S.A \| Japan | intron | 5677285 | France |
| *CG1304* | 1.656842 | U.S.A \| Japan | flank2kb | 2387338 | Japan |
| *Mal-A3* | 1.215228 | U.S.A \| Japan | flank2kb | 5257029 | France |
| *Tsf1* | 1.261203 | U.S.A \| Japan | intron | 1957700 | Jap |

**Table S8.** Geographical location of isofemale lines. *D. suzukii* flies were sampled in 3 countries (Japan, U.S.A and France) with their location and invasive status. Line name is indicated with bold type for the line use in molecular analysis.

| Location | Coordinates | Status | Lines |
| --- | --- | --- | --- |
| Sapporo (Hokkaido, Japan) | 43° 3' 43.545"N 141° 21' 15.754" E | Native | S11, S20, S21, S24, **S29** |
| Tokyo (Honshu, Japan) | 35° 41' 22.155" N 139° 41' 30.143" E | Native | T3, T11, T18 |
| Watsonville (California, U.S.A) | 36°54'51.8"N 121°45'27.7"W | Invasive | W106, W112, W113, **W120**, W122, W127 |
| Dayton (Oregon, U.S.A) | 45° 13' 14.422" N 123° 4' 34.368" E | Invasive | Sok1, Sok28, Sok58 |
| Paris (France) | 48° 51' 23.81" N 2° 21' 7.998" E | Invasive | L2, L6, L7, L21, L22, L26 |
| Montpellier (France) | 43° 36' 38.768" N 3° 52' 36.177" E | Invasive | MT15, MT20, MT25, **MT47** |

**Table S9.**
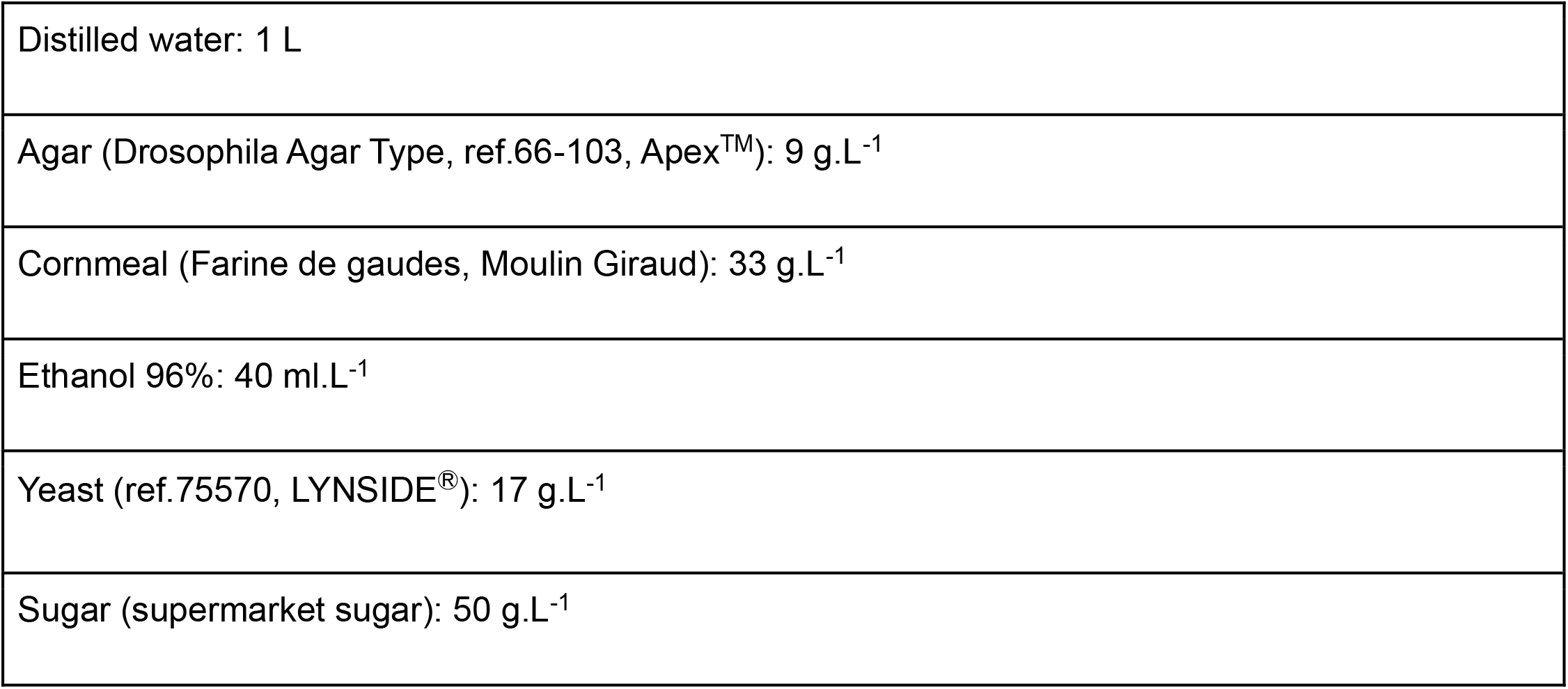
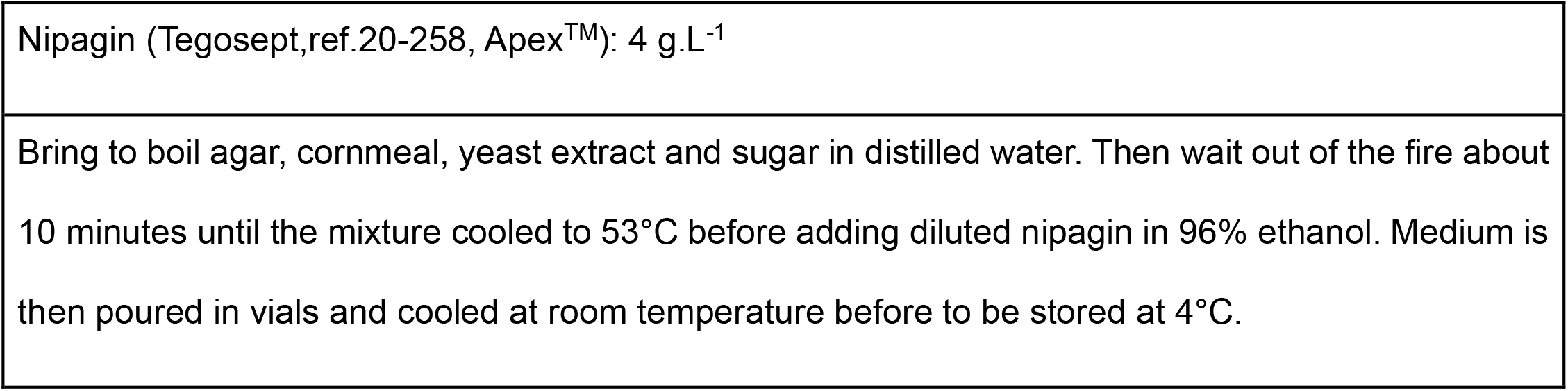
Recipe of diet medium modified from Dalton *et al*., 2011.

## Notes

ftp://pbil.univ-lyon1.fr/pub/datasets/Marin2020/surv_suzukii_marin2020/surv_suzukii_marin2020.txt

### Competing Interest Statement

The authors have declared no competing interest.

